# Layer 1 NDNF interneurons form distinct subpopulations with opposite activation patterns during sleep in freely behaving mice

**DOI:** 10.1101/2025.01.10.632405

**Authors:** Aurélie Brécier, Gaëlle Mailhos, Przemyslaw Jarzebowski, Yuqi Li, Ole Paulsen, Audrey Y. Hay

## Abstract

Non-rapid eye movement (NREM) sleep facilitates memory consolidation by transferring information from the hippocampus to the neocortex. This transfer is thought to occur primarily when hippocampal sharp-wave ripples (SWRs) and thalamocortical spindles are synchronized. However, the mechanisms underlying this synchronization remain unknown. In this study, we investigated the role of cortical layer 1 NDNF-expressing (L1 NDNF) interneurons in gating information transfer during SWR-spindle synchronization in NREM sleep. Using simultaneous cell-specific calcium imaging with a head-mounted microscope and local field potential recordings in freely moving mice, we compared the activity of L1 NDNF and L2/3 neurons across vigilance states and during NREM-specific oscillations. Our findings reveal that L1 NDNF neurons form three distinct populations, assembling into cell networks tuned to specific sleep stages. REM active L1 NDNF and L2/3 neurons exhibit opposite activation patterns during spindles. While L2/3 cells are mostly inactive during SWR, NREM, and REM active L1 NDNF cells inhibit the network upon SWR onset depending on their coupling with spindles. L1 NDNF neurons mediate slow inhibition primarily via GABA_B_ receptors. Systemic application of a GABA_B_ receptor antagonist resulted in decreased neuronal coupling of pyramidal cells but did not change the inhibitory responses during SWRs. Overall, these findings highlight the potential role of L1 NDNF neuron-mediated inhibition in the response to synchronized sleep oscillations, with potential implications for memory consolidation.

**Significance Statement:** This study highlights how a specific type of inhibitory neuron, L1 NDNF-expressing interneurons, is activated during deep sleep. We identified three different subpopulations of L1 NDNF neurons based on their activity during vigilance states. These subpopulations show specific and complementary activation patterns during two key sleep oscillations-sharp-wave ripples (SWRs) and spindles-which are crucial for transferring memories from short-term to long-term storage. This research provides new insights into how brain oscillations synchronize and how this phenomenon correlates with local neuronal activity, with potential implications for understanding learning and memory mechanisms.

## Introduction

Sleep, and in particular non-rapid-eye movement (NREM) sleep, plays an important role in transforming daily experiences into lasting memories ^1,2^. This memory consolidation process is supported by a dialogue between the hippocampus, which initially encodes the information, and the neocortex, which holds the long-term stable memories ^3^. This dialogue is thought to occur when hippocampal sharp-wave ripple (SWR) oscillations (120-200 Hz) are coordinated with neocortical slow oscillations (0.5 – 2 Hz) and thalamocortical spindles (10-16 Hz) ^4–9^. Indeed, the coupling of these oscillations increases during NREM sleep following learning and promoting the coupling between SWRs, slow oscillations and spindles improves memory consolidation ^9,10^. However, how this coordination is orchestrated remains unclear.

Layer (L)1 of the neocortex is a major neocortical hub where dendritic tufts from L2/3 and L5 pyramidal neurons integrate inputs from multiple subcortical regions, in particular from the thalamus, and from other neocortical areas ^11,12^. Integration of these inputs is controlled by local inhibitory neurons that also receive long-range and local inputs and target the tufted dendritic branches of pyramidal neurons. Among these, a subclass of L1 interneurons, the neurogliaform cells, or L1 neuron-derived neurotrophic factor (NDNF) interneurons is of major interest ^13,14^. L1 NDNF neurons are targeted by corticocortical and subcortical inputs ^14,15^ and exert a powerful control over pyramidal cell activity across all cortical layers through both fast (GABA_A_) and slow (GABA_B_) inhibition ^16–19^. Additionally, recent research has linked their activity with memory performance in mice ^20,21^. Although a role for L1 NDNF interneurons in generating Down states has been identified ^15^, it remains uncertain whether they also contribute to the integration or synchronization of SWRs and spindles.

Here we performed simultaneous cell-specific one-photon calcium imaging, pharmacological manipulations, and local field potential (LFP) recordings in two neocortical areas and the hippocampus of freely moving, naturally sleeping mice. We found that each L1 NDNF neuron preferentially activates during a specific brain state — wakefulness, rapid-eye movement (REM) sleep, or non-REM (NREM) sleep — and this state-specific activity remains stable over multiple days. Moreover, L1 NDNF neurons with similar activity profiles form assemblies with each other, suggesting coordinated network-level organization. These three subpopulations also differentially link to sleep oscillations, with L1 REM cells being particularly recruited when spindles are simultaneously detected in different brain regions. L1 NDNF neuron subpopulations also differentially and complementary responded to SWR-spindles synchronization. Finally, blocking GABA_B_ receptors (GABA_B_R), the main receptor family activated by L1 NDNF neurons, decreased the neuronal coupling of pyramidal neurons while maintaining the response of the network to sleep oscillations. Taken together, this study suggests a prominent role for L1 NDNF neuron-mediated inhibition in 1) controlling the cortical activity during and around spindles, and 2) dynamically inhibiting the network during the coupling of hippocampal SWRs and neocortical spindles, which could affect memory consolidation.

## Results

### L1 NDNF population consists of Wake active, NREM active and REM active subpopulations

We combined one-photon calcium imaging, using Miniscope V4, with LFP recordings of the contralateral prefrontal cortex (PFC), the primary somatosensory cortex (S1), and hippocampal CA1 to monitor the activity of cortical neurons in naturally sleeping mice (Fig. 1A-C). First, we injected a non-Cre-dependent calcium indicator-GCamp6f-in the L2/3 of the dorsal neocortex of NDNF-cre mice (*N* = 3 mice) (Fig. 1D). We then simultaneously recorded LFPs and the activity of the putative L2/3 pyramidal cells during the sleep-wake cycle (Fig. 1E-F). Vigilance states were scored over time using raw and filtered LFPs, and electromyographic (EMG) recordings (Fig. 1C). We first confirmed that our recordings with Miniscope did not affect the sleep quality of the mice by recording the same animals with and without the Miniscope (Fig. S1A-C). Then, calcium transients in identified neurons were extracted and synchronized with the vigilance states. The normalized area under the curve of the calcium signal of each neuron was computed for each episode. We first quantified the averaged activity in active wake (AW), quiet wake (QW), NREM, and REM sleep. We found that L2/3 neurons had higher activity at the population level during wake and REM sleep compared to NREM sleep (Fig. 1G, see also S2A and Table S1 for statistical analysis). We then asked if the averaged activity reflects the behavior of individual neurons. For that, we represented the activity of each cell on a uniform manifold approximation and projection (UMAP) (see Methods) reflecting the preferential activity of each neuron during AW, QW, NREM, and REM sleep. We found that L2/3 cells were homogenously spread on the UMAP, and that most cells are active during both AW, QW, and REM sleep (Fig. 1H).

**Fig. 1.**
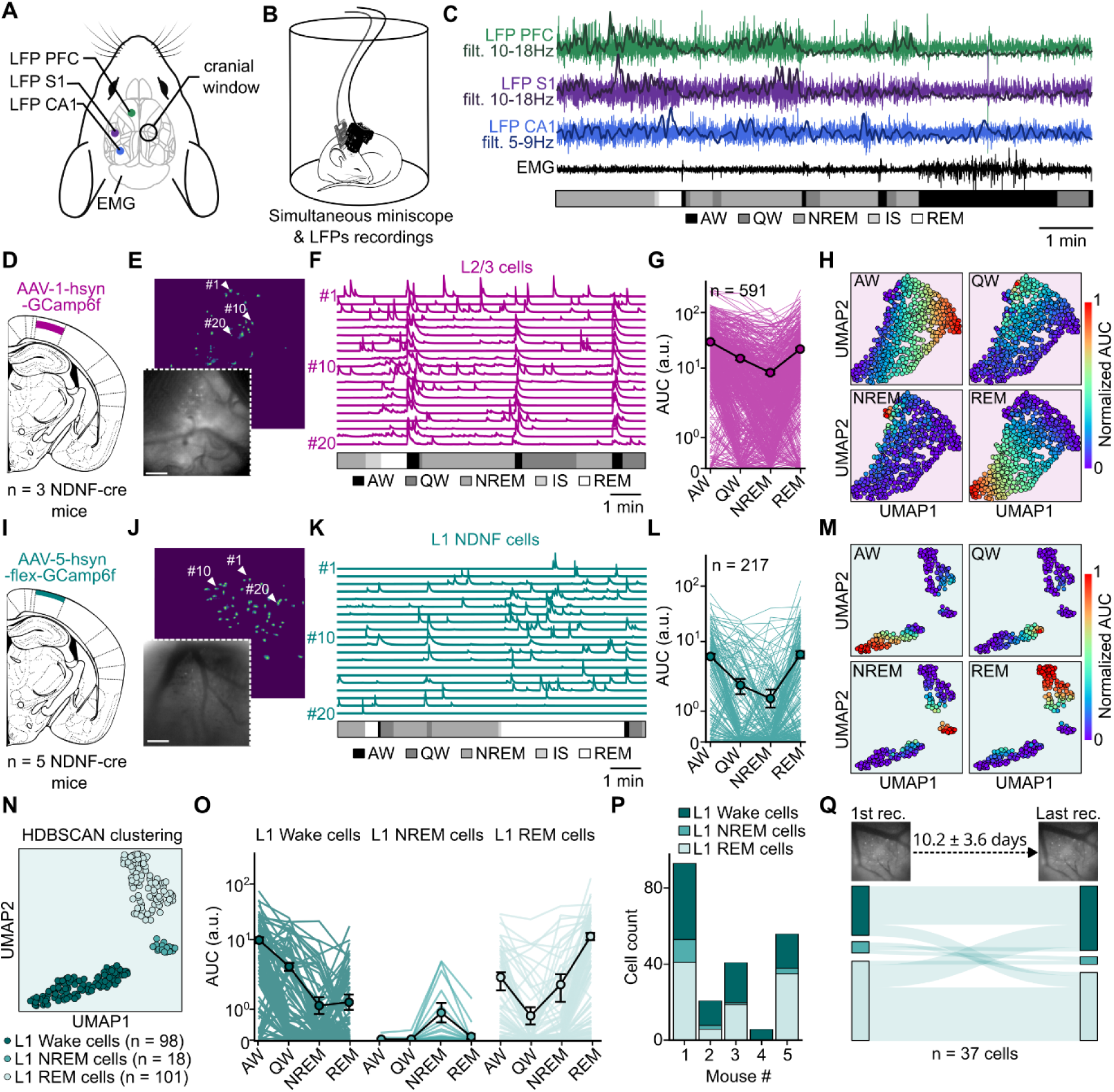
L1 NDNF population consists of Wake active, NREM active and REM active subpopulations. (**A**) Schematic of the LFP and Miniscope implantation. (**B**) Representation of a sleeping mouse wearing a Miniscope and a connected LFP/EMG headstage. (**C**) Top, raw LFPs of PFC, S1, CA1, with their respective filtered (filt.) LFPs superimposed and EMG. Bottom, the hypnogram of the recording example. (**D**) Viral injection site in layer 2/3 with a non Cre-dependent GCamp6f construct (AAV-1-hsyn-GCamp6f) in NDNF-Cre mice. (**E**) Field of view under the Miniscope in a mouse expressing GCamp6f in L2/3 neurons and associated spatial footprint. (**F**) Calcium traces of 20 L2/3 neurons in E according to the vigilance states. Scale bar 200 µm. (**G**) Average area under the curve (AUC) of the calcium transients of unique L2/3 neurons recorded in all sleep states (P < 0. 001 for all comparisons, except P_REM-QW_ > 0.05). (**H**) UMAP based on the activity of L2/3 cells in G in each vigilance state. (**I**) Viral injection site in layer 1 with a Cre-dependent GCamp6f construct (AAV-5-hsyn-flex-GCamp6f) in NDNF-Cre mice. (**J**) Field of view under the Miniscope in a mouse expressing GCamp6f in L1 NDNF neurons and associated spatial footprint. Scale bar 200 µm. (**K**) Calcium traces of 20 L1 NDNF neurons in J according to the vigilance states. (**L**) Average AUC of the calcium transients of unique L1 NDNF neurons recorded in all vigilance states (P < 0.001 for all comparisons, except P_AW-REM_, P_NREM-QW_ > 0.05). (**M**) UMAP based on the activity of L1 NDNF cells in L in each vigilance state. (**N**) Results of the HDBSCAN clustering performed on L1 NDNF UMAP. Three distinct clusters were identified. (**O**) Average AUC of the calcium transients of each L1 NDNF neurons clusters according to sleep states (For L1 Wake cells: P < 0.001 for all comparisons, except P_NREM-QW_ < 0.05 and P_NREM-REM_, P_NREM-QW_ > 0.05; For L1 NREM cells: P_AW-NREM_ and P_NREM-QW_ < 0.01; For L1 REM cells: P < 0.001 for all comparisons, except P_AW-QW_ < 0.05 and P_NREM-AW_, P_NREM-QW_ > 0.05). (**P**) Proportion of L1 Wake, NREM and REM cells for each recorded mouse. (**Q**) Classification of L1 NDNF neurons recorded over more than 2 recording sessions according to their cluster during their first and last recording session. Generalized linear mixed-effects model (G, L, and O). See Table S1 for statistics. Data are shown as means ± SEM.

We next performed the same experiment on NDNF-cre mice injected with a Cre-dependent AAV containing GCamp6f in order to monitor the activity of L1 NDNF cells specifically (*N* = 5 mice) (Fig. 1I-K, and S1D). Our results indicate that at the population level, L1 NDNF cells present an increased activity during active wake and REM sleep, similarly to L2/3 cells (Fig. 1L, see also S2B). However, the representation of L1 NDNF cell activity on a UMAP (using the same parameters as in H), revealed three distinct clusters of neurons: L1 NDNF cells mostly active during AW and QW, L1 NDNF cells active during NREM and L1 NDNF cells active during REM sleep (Fig. 1M, S2C-D). The unsupervised clustering method confirmed the presence of these three clusters of neurons that we labeled L1 Wake cells, L1 NREM cells, and L1 REM cells, respectively (Fig. 1N). The averaged activity of these three clusters of neurons confirmed their respective preference for Wake, NREM, or REM sleep, respectively (Fig. 1O, S2E-F). We then controlled the presence of the three subpopulations of neurons in each of our recorded mice (Fig. 1P, S2G-H) and found that all mice but one presented L1 NDNF neurons from all three subtypes. The remaining mouse presented only L1 Wake cells, which might be due to the small sample of neurons recorded from this mouse (< 10 cells). Surprisingly, we found that on average neurons belonging to the same cluster were more distant from each other compared to cells from different clusters. This was particularly prominent for L1 REM cells (Fig. S2I-J). Lastly, we investigated whether sleep stage selectivity was preserved over days. We compared the cluster identity on the first and last days of recording (10.2 days on average, ranging from 1 to 48 days) for each cross-registered neuron. Data revealed that the assignment of a L1 NDNF neuron to a cluster remained stable for 71 % of the neurons (Fig. 1Q).

These findings reveal the existence of distinct subpopulations of L1 NDNF neurons with consistent sleep-stage-specific firing patterns, contrasting with the absence of selectivity observed in L2/3 neurons. Furthermore, the analysis uncovered unique populations of Wake-, NREM-, and REM-active L1 NDNF neurons that were not previously identified using traditional averaged activity assessments ^19,22^.

### Stage-selective neurons form cell assemblies

Next, we investigated whether L1 NDNF neurons belonging to the same cluster take part in similar neuronal assemblies. We first calculated the correlation coefficient for each pair of neurons according to their cluster identity during the entire recording (wake and sleep combined). We found that L1 NDNF neurons selective for either Wake or REM sleep have a higher probability of firing with neurons from the same cluster compared to neurons from different clusters (Fig. 2A-C). We next compared how subpopulations of neurons were correlated to themselves during the vigilance state. Our results show that the activity of neurons belonging to the same cluster exhibited highly correlated activity during both active and quiet wake (Fig. S3A-B). L1 REM neurons exhibited particularly strong pairwise correlations during wakefulness. Overall, this indicates 1) that the correlation strength is not influenced by the average activity, and 2) that wakefulness strengthens coordinated activity within neuronal subpopulations.

**Fig. 2.**
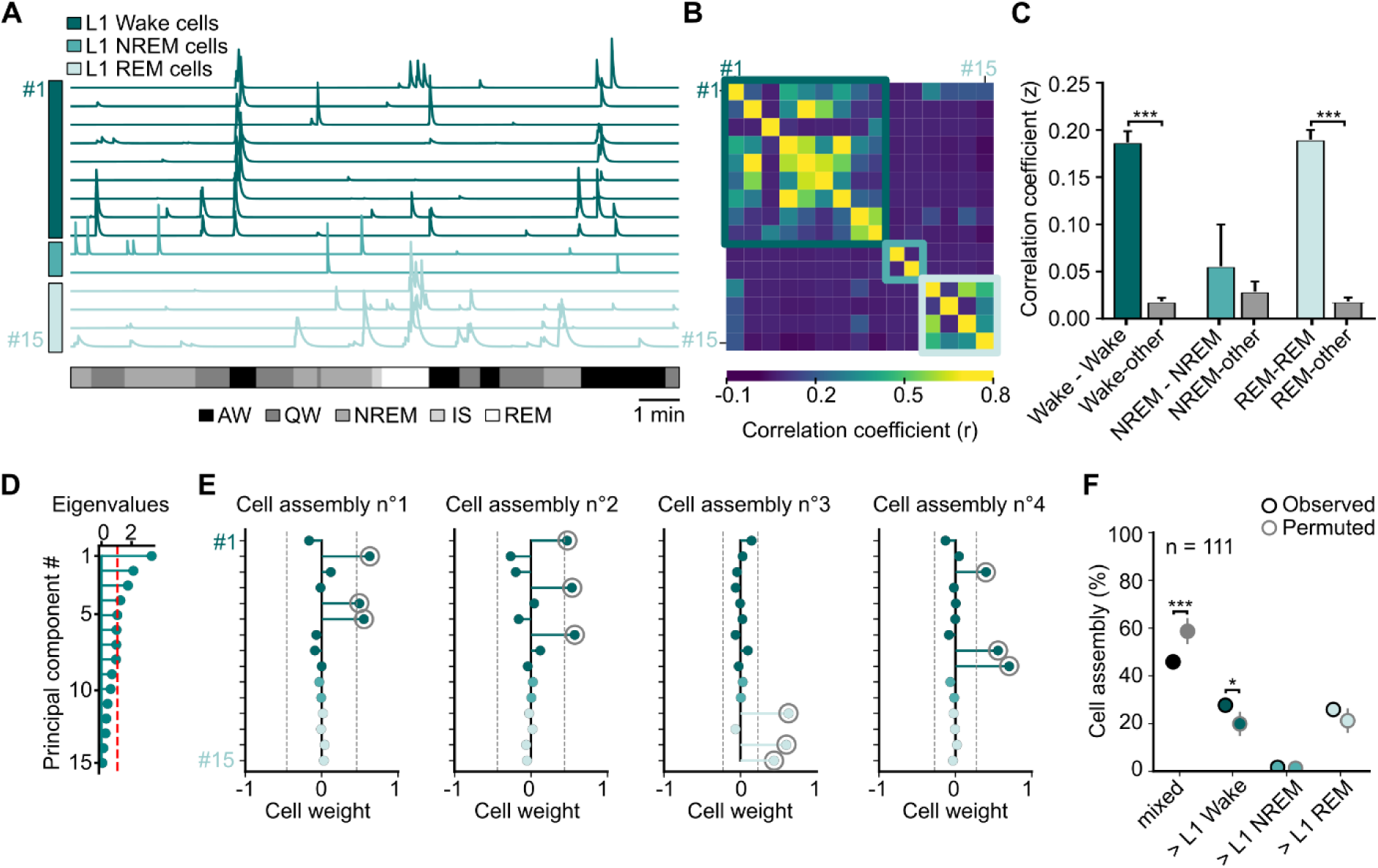
L1 NDNF neurons from the same subpopulation fire together. (**A**) Recording example of 15 L1 NDNF neurons across vigilance states. Neurons were ranked according to their cluster identity. (**B**) Heatmap of the correlation coefficient (*r*) between each calcium transients displayed in A. Colored squares define L1 Wake, NREM and REM cells. (**C**) Average z-transform correlation coefficients (z) between L1 Wake cells, between L1 Wake cells and the rest of the population, between NREM cells, between NREM cells and the rest of the population, between L1 REM cells, between L1 REM cells and the rest of the population (Wake-Wake, n = 615 pairs; Wake-other, n = 1309 pairs; NREM-NREM, n = 14 pairs; NREM-other, n = 386 pairs; REM-REM, n = 712 pairs; REM-other, n = 1291 pairs). (**D**) Eigenvalue distribution from the example in Fig. 2A. Red dotted line represent the Marcenko-pastur threshold that determine significant principal components (i.e. cell assemblies). In this example, 5 putative cell assemblies were detected. (**E**) Neurons weight in each detected cell assemblies in the example in A. Dotted grey lines represent the threshold for a neuron to cross to significantly participate in the cell assembly. Here, one cell assembly was removed because only 1 neuron crossed the threshold. Note how cell assemblies are mostly constituted of neurons belonging to the same cluster. (**F**) Proportion of cell assembly observed containing a majority of L1 Wake cells (> L1 Wake), L1 NREM cells (> L1 NREM), L1 REM cells (> L1 REM) or a mixed population of cells (mixed) compared to the expected proportion given the distribution of cell subpopulations (n = 5000 permutations). *P < 0.05, ***P < 0.001. One-way anova (C) and permutation tests (F). See Table S1 for statistics. Data are shown as means ± SEM.

We then wondered if neurons from the same cluster are organized into cell assemblies. We computed a principal component analysis on calcium transients followed by an independent component analysis (see Methods) to identify significant cell assemblies and their respective cell members for each recording (Fig. 2D-E, S3C-D). We then classified cell assemblies according to the cluster identity of their cell members. Our results indicate that given the proportion of L1 Wake, NREM, and REM cells in each recording, the probability of getting cell assemblies composed of multiple cell subtypes (mixed) was lower in our sample than expected by chance (Fig. 2F). This suggests that cells from the same cluster tend to form cell assembly, further reinforcing the idea of distinct subpopulations in L1 NDNF cells.

### L1 NDNF and L2/3 neurons show opposite activity patterns around spindles

During NREM sleep, thalamocortical spindles can be recorded locally in most neocortical areas, but are prominently observed in the PFC and S1. Spindles are thought to play a role in memory consolidation by promoting synaptic plasticity locally. Because L1 NDNF neurons could affect plasticity processes specifically on distal dendritic tufts, we investigated the activity of L1 NDNF subpopulations and L2/3 neurons around spindles. In our recordings, we defined 3 types of spindles: those detected in PFC but not S1 (PFC spindles), those detected in S1 but not in PFC (S1 spindles), and those recorded simultaneously in S1 and PFC (S1&PFC spindles) (Fig. 3A). Spindles detected in both S1 and PFC comprised about 15 % of all detected spindles (Fig. 3B and S4A). Of note, the probability of recording spindles in both S1 and PFC was significantly higher than chance level (Fig. 3C), suggesting that distinct mechanisms underlie the generation of putative global versus local spindles. S1&PFC spindles lasted longer and were equally likely to first be detected in either S1 or PFC (Fig. S4B-D).

**Fig. 3.**
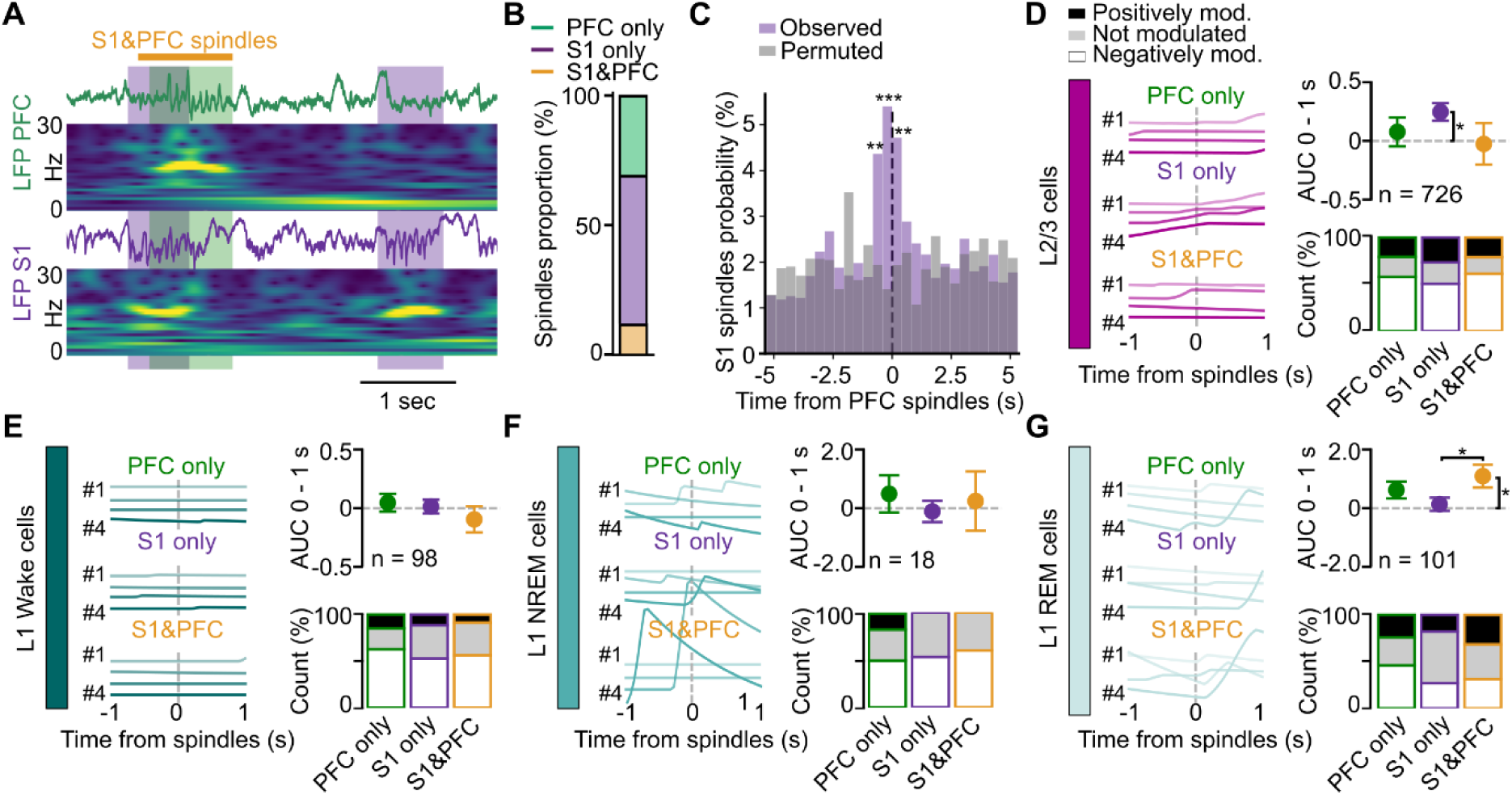
The activity of L1 NDNF subpopulations and L2/3 cells differentially correlate with spindles depending on spindle location. (**A**) Example recording of raw LFPs from PFC, S1 and their respective spectrogram from 0 to 30 Hz. Green and purple shaded areas represent detected spindle oscillations in PFC and S1, respectively. Spindles occurring simultaneously in both areas are defined as S1&PFC spindles. (**B**) Average proportion of spindles detected depending on their locations (*N* = 8 mice). (**C**) Observed occurrence probability of S1 spindles around PFC spindles compared to permuted probability (n = 5000 permutations). (**D**) Example of one L2/3 neuron calcium activity 1 second before and after detected spindle onset (time 0) according to spindle locations (left). Average baselined AUC (a.u.) of L2/3 neurons after spindle onset (top right). Proportion of unmodulated, positively modulated and negatively modulated cells by spindles as a function of spindles location (bottom right). (**E**) Same as in D for the L1 Wake. (**F**) Same as in D for the L1 NREM. (**G**) Same as in D for the L1 REM neurons. *P < 0.05, **P < 0.01, ***P < 0.001. Generalized linear mixed-effects model (D, E, F and G). See Table S1 for statistics. Data are shown as means ± SEM.

Since our calcium imaging recordings were performed contralaterally to the LFPs recordings (due to space restriction on the skull), we first controlled that the majority of spindles detected in one hemisphere were also present in the contralateral one. Our measurements revealed that more than 90% of PFC spindles occur in both cortices (Fig. S4E-G). Similarly, a majority of S1 spindles (more than 70%) were recorded simultaneously in both cortices (Fig. S4H-J). We therefore evaluated the neuronal activity in the dorsal neocortex during and around these 3 types of spindles, contralaterally to where spindles were detected. L2/3 neurons activity increased during S1 spindles but did not significantly differ between PFC, S1, and S1&PFC spindles (Fig. 3D). We next investigated the activity of L1 Wake, NREM, and REM cells around spindles. On average, both L1 Wake and NREM cell activity remained unchanged during spindles, independently of spindles location (Fig. 3E-F). In contrast, L1 REM cell activity was significantly increased during S1&PFC spindles specifically (Fig. 3G). These findings suggest that putative global spindles particularly drive L1 REM neuron activity in comparison to isolated spindles. Interestingly, the entire population of L1 NDNF cells tends to behave on average as L1 REM cells, with higher activity during S1&PFC spindles, but also during PFC spindles (Fig. S4K). Of note, we were not able to identify any significant L1 NDNF or L2/3 cell assemblies reactivation around spindles (Fig. S4L-M). Taken together, our results suggest that L1 NDNF neurons could inhibit L2/3 neurons during S1&PFC spindles while, during S1 spindles, decreased activity of L1 NDNF neurons could permit L2/3 neurons to increase their activity. Moreover, our results revealed that L1 NDNF neurons with higher activity during REM sleep are the ones that strongly activate during S1&PFC spindles, which could have interesting implications for memory consolidation as discussed in the Discussion section.

### L1 NREM and REM cells activity increases after CA1 SWR and is modulated by the coupling of SWR to spindles

SWR-spindle coupling is a hallmark of memory consolidation. Since neuronal activity is differentially modulated by spindle synchrony, we next hypothesized that the coupling of hippocampal SWRs with spindles could also be associated with specific neuronal network activity. We quantified the percentage of spindles with at least one SWR recorded in CA1 (within 0.5 second before or during the spindle (Fig. 4A). PFC and S1&PFC spindles were significantly more coupled to SWR than S1 spindles (Fig. S5A), consistent with the reported role of active PFC spindle – CA1 SWR coupling in memory consolidation ^23,24^. We controlled that SWRs co-occur in both hemispheres and found that 90% of the CA1 SWRs detected in one hemisphere were observed in the other one (Fig. S5B-D).

**Fig. 4.**
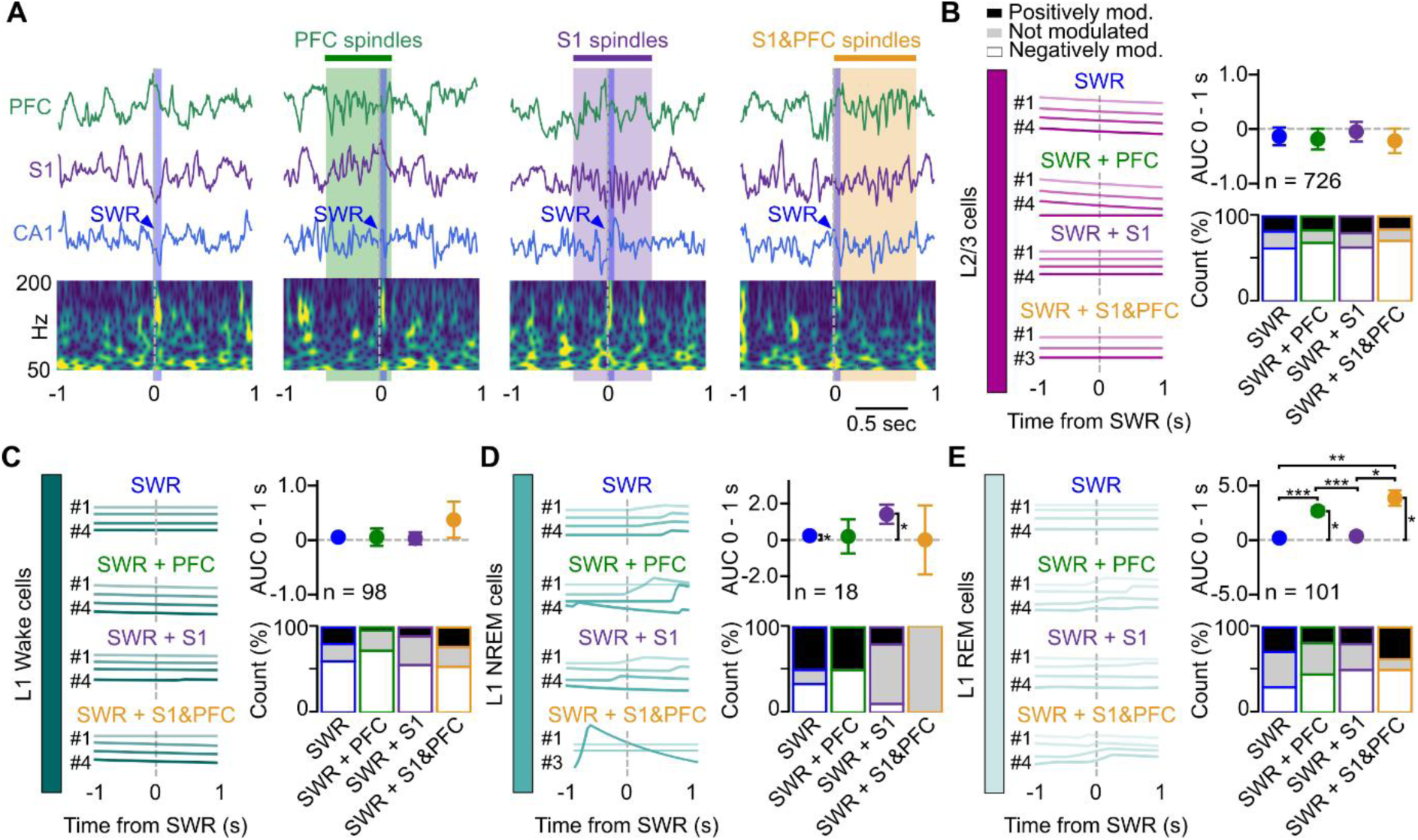
SWR - spindle coupling is correlated with increased activity of L1 NREM and REM cells. (**A**) Example of LFP recordings of PFC, S1 and CA1 around detected SWR (blue shaded area). SWR were considered coupled with a PFC, S1 or S1&PFC spindles (green, purple or orange shaded area, respectively) if the SWR occurred 500 ms before or during the spindle. (**B**) Example of one L2/3 neuron calcium activity 1 second before and after the onset of uncoupled SWR, SWR coupled with PFC spindles, with S1 spindles or S1&PFC spindles (left). Average baselined AUC (a.u.) of L2/3 neurons after SWR onset (top right). Proportion of unmodulated, positively modulated and negatively modulated cells by SWR as a function of its coupling (bottom right). (**C**) Same as in B for the L1 Wake. (**D**) Same as in B for the L1 NREM. (**E**) Same as in B for the L1 REM neurons. *P < 0.05, **P < 0.01, ***P < 0.001. Generalized linear mixed-effects model (B, C, D, and E). See Table S1 for statistics. Data are shown as means ± SEM.

In order to investigate whether oscillation coupling affects neuronal activity, we compared the activity of L2/3 neurons and L1 NDNF around SWR not coupled with spindles, coupled with PFC spindles, coupled with S1 spindles and coupled with S1&PFC spindles. Our data indicate that the occurrence of spindles coupled with SWR did not, on average, influence the activity of L2/3 neurons after SWR onset (Fig. 4B). More L2/3 cells than expected by chance were negatively modulated by SWR, regardless of their coupling (Observed proportion: 64.7 %; Random proportion: 51.2 %; p < 0.001). At the cell assembly level, we uncovered a reduction of the activity of L2/3 cell assemblies prior to SWR (Fig. S5E), consistently with the literature ^25^. The activity of L1 NDNF subpopulations relative to SWR showed contrasted results. The activity of L1 Wake cells was unaffected by SWR whether they were or not coupled to spindles, while L1 NREM cells activity was mildly increased after uncoupled SWRs or SWRs coupled with S1 spindles (Fig. 4D). In contrast, L1 REM cells activity was strongly impacted by the occurrence of SWR. When coupled with a PFC or S1&PFC spindles, L1 REM cell activity was increased in comparison to the activity recorded during an uncoupled SWR or a SWR coupled with S1 spindles (Fig. 4E). L1 NDNF cells, at the population level, behave similarly to L1 REM cells while the activity remains constant at the cell assembly level (Fig. S5F-G). These findings highlight a strong activation of L1 REM neurons around SWR synchronized with spindles occurring in the PFC, which mirrors the mild inhibition seen in L2/3 neurons. In contrast, L1 NREM cells were activated during uncoupled SWR or SWR coupled with S1 spindles, thus suggesting a potential complementary role of L1 REM and L1 NREM in promoting SWR induced inhibition of the neocortex.

### GABA_B_R-mediated inhibition modifies L2/3 neuronal coupling

L1 NDNF neurons can control L2/3 neurons excitability through GABA_B_R activation ^19,26^. Thus, we tested the impact of blocking GABA_B_R-mediated inhibition on neuron activity across the sleep-wake cycle. CGP55,845, a GABA_B_R antagonist, was injected intraperitoneally and we recorded the neuronal activity before and one hour after the injection (Fig. 5A and B). We first controlled that CGP55,845 injection did not have any impact on sleep quality. Our results did not reveal any change in the percentage of time spent in each vigilance state nor in episode duration upon CGP55,845 injection (Fig. S6A-B). We found that L2/3 neurons average activity was not changed during vigilance states after the CGP55,845 injection (Fig. 5C).

**Fig. 5.**
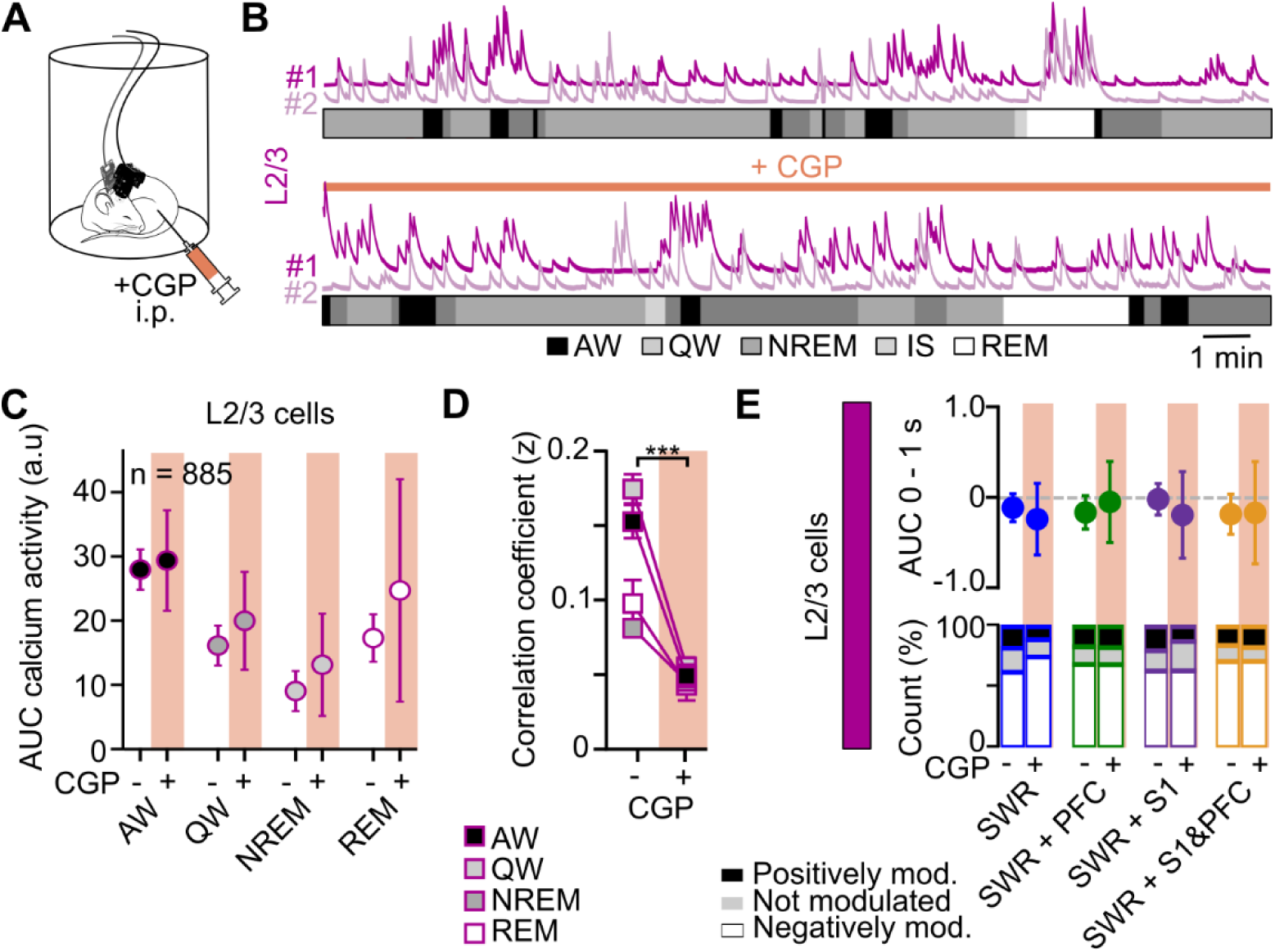
L2/3 neuronal coupling is GABA_B_R-dependent. (**A**) Implanted mice received an i.p. injection of the selective GABA_B_R blocker CGP55,845. (**B**) Two examples of L2/3 neuron activity before (top) and after (bottom) CGP55,845 injection. (**C**) Average AUC of L2/3 neuron calcium activity before and after CGP55,845 injection according to vigilance states. (**D**) Evolution of the correlation coefficient (z) before and after CGP55,845 injection during all vigilance states (n = 2568 pairs). (**E**) Average baselined AUC (a.u.) of L2/3 neurons after SWR onset (top). Proportion of unmodulated, positively modulated and negatively modulated cells by SWR as a function of its coupling and CGP55,845 injection (bottom). ***P < 0.001. Generalized linear mixed-effects model (C and E) and two-way anova (D). See Table S1 for statistics. Data are shown as means ± SEM.

L1 NDNF neurons could affect L2/3 neuronal coupling through synchronized inhibition of pyramidal dendritic tuft mediated by GABA_B_R activation. Consistent with this hypothesis, data revealed that the activity of L2/3 neurons was less synchronized after CGP55,845 injection (Fig. 5B and D). Interestingly, L1 NDNF cell activity, coupling, and vigilance state preference remained constant under CGP55,845 (Fig. S6C-E), suggesting that GABA_B_Rs have a limited impact on the activity regulation of L1 NDNF neurons.

We then wondered if this decorrelation of L2/3 cells could have an impact on spindle response. On average, CGP55,845 did not change the spindle rate (Fig. S6F), nor the activity of L2/3 and L1 NDNF cells upon PFC, S1 and S1&PFC spindles onset (Fig. S6G-J). We next investigated whether the GABA_B_R blocker CGP55,845 could change SWR-spindle synchrony and related neuronal activity. SWR rate as well as spindles-SWR coupling remained constant under CGP55,845 condition (Fig. S6K-L). Similarly, CGP55,845 had no effect on L2/3 cells and L1 NDNF neurons activity during hippocampal SWRs, independently of its coupling (Fig. 5E and S6M-O). Overall, these results suggest that GABA_B_Rs-mediated slow inhibition, possibly generated by neurogliaform cells, could have a role in synchronizing the activity of L2/3 neurons, but do not appear to change the activity pattern of L2/3 cells during oscillation synchronization.

## Discussion

We performed simultaneous recordings in CA1, S1 and PFC in combination with calcium imaging of a hitherto underappreciated class of interneurons in freely-moving mice to unveil their function in sleep oscillations. Our results revealed that L1 NDNF neurons form subpopulations strongly tuned to specific vigilance states. Moreover, we showed their differential activation depending on spindle location and their complementary inhibitory role during spindle-SWR coupling. Finally, we found that GABA_B_R-mediated inhibition reduces the coordinated activation of pyramidal cell without impacting sleep structure, or neuron’s response to sleep oscillations.

### L1 NDNF neurons are strongly vigilance-state tuned

L1 NDNF cells like other cortical interneurons ^27,28^ show variable activity levels depending on the vigilance state. Our study and others ^19,22^ revealed that L1 NDNF neurons are on average more active during wakefulness and REM sleep compared to NREM sleep. However, analysis of individual cell activity revealed particular subpopulations preferentially and reliably active during Wake, NREM, or REM sleep. REM and wake are characterized by high acetylcholine release in the neocortex while cholinergic tone is low during NREM sleep ^29^. L1 NDNF neurons are sensitive to acetylcholine ^30^ but L1 interneurons show variability in the expression of nicotinic receptors ^31,32^. It would be interesting to test whether L1 NDNF neurons differ in their nicotinic receptor expression and if this could drive their selectivity.

Previous studies highlighted heterogeneity within the L1 NDNF neuron population. In particular, late-spiking and regular-spiking phenotypes were identified forming two distinct cell subpopulations ^13,14^. The latter profile was related to the specific expression of Neuropeptide Y in the somatosensory cortex ^14^, while it did not correlate to any genetic targets in the auditory cortex ^13^. Because of their persistent firing proprieties and their late spiking phenotype, L1 NDNF neurons have been described as a cell type optimized for slow signals ^13,33^ thus it would be interesting to test if L1 NREM are of the late spiking type. A recent study also identified subpopulations of L1 cells with distinct transcriptomic profiles that exhibited differential responses to state modulation ^34^. Specifically, two subpopulations (Lamp5-Npy cells and Lamp5-Plcj2-Dock5 cells) were found to be more active during active wakefulness, while a third subpopulation (Lamp5-Lsp1 cells) showed increased activity during synchronized states (putative quiet wake and NREM sleep). Further investigations using in vivo patch-clamp recordings for example could be helpful to understand if the identified L1 Wake, NREM, and REM cells match the different spiking phenotypes and/or genetic profiles. It would also be of interest to investigate whether these three clusters of layer 1 NDNF neurons are conserved across other cortical areas, which could provide insights into their broader functional relevance within cortical circuits.

### Spindle synchrony differentially impacts L1 NDNF cells

Although spindles are primarily local events, global spindles have also been observed in both humans ^35,36^ and mice ^37^. While the particular role of global spindles in memory consolidation remains to be understood, there is evidence to suggest that spindle synchrony is regulated by cortico-cortical and corticothalamic projections ^38^ as well as glutamatergic neurotransmission ^39^. In line with these studies, our work revealed the involvement of distinct cortical networks depending on the spatial organization of spindles. In particular, putative global spindles were associated with strong responses of L1 NDNF neurons. These findings suggest a powerful inhibition of the apical dendrites of pyramidal cells during synchronized spindles, potentially impacting plasticity outcomes. On the other hand, somatosensory spindles detected contralaterally were associated with a moderate inhibition of L1 NDNF neurons and a robust activation of L2/3 neurons. While L1 NDNF and L2/3 neurons display complementary activity, blocking L1 NDNF slow inhibition did not alter the observed response. However, we cannot exclude that L1 NDNF neurons mediate feed-forward inhibition during global spindles through GABA_A_Rs onto pyramidal cells.

Surprisingly, REM-active L1 NDNF neurons, while representing less than half of the population of L1 NDNF cells, drive the activation observed during putative global spindles. These neurons were also the most correlated ones during wakefulness compared to NREM and REM neurons. This suggests specific reactivation of the L1 interneurons recruited during REM sleep and global spindles, which could speculatively be of particular importance for memory consolidation.

### L1 NDNF neuron activity is modulated by SWR-spindle coupling

Recent findings suggest that the coupling between SWR and spindles is crucial for memory consolidation. More specifically, memory performance on a hippocampus-dependent task was improved by artificially increasing this synchronization ^10^. Others argue that spindles set a timeframe for SWR to occur ^6,9^. Interestingly, our study reveals that the activity of L2/3 pyramidal cells is not influenced by the presence of a coupling between SWRs and spindles, which is not the case for L1 NDNF neurons. Our findings uncover a complementary inhibitory role of L1 NDNF cells active upon SWR. L1 NDNF cells active during REM sleep reactivate during SWR coupled with PFC spindles while L1 NDNF neurons active during NREM only respond to uncoupled SWR and SWR coupled with S1 spindles. These results could indicate that L1 NDNF subpopulations are differentially connected to distal cortical areas. It could also explain the absence of activity change in pyramidal cells depending on the SWR-spindle synchrony. Last, it suggests a particular mechanism linking the activity during REM and NREM sleep, with a potential role for synaptic plasticity.

Similar to what has been described in the past, we observed that cortical activity around hippocampal SWRs is mostly inhibitory ^25,40^. It has been reported that pyramidal neurons tend to reduce their activity prior to SWRs occurring during wake, a phenomenon also visible in our sleep recordings. Similarly to what has been observed during wakefulness ^25^, sleep SWRs induce a modest inhibition of pyramidal cells. However, we did not observe a reduced activity of L1 interneurons around SWR occurrence. This difference further emphasizes the distinct effects of sleep and awake SWRs on cortical activity.

### Slow inhibition increases pyramidal cells synchrony but does not contribute to oscillation coupling

It is now well established that a major, but not exclusive, source of GABA_B_R-mediated inhibition in the cortex is mediated by L1 NDNF neurons ^14,16–18,20^. In addition, Hay et al. have demonstrated that L1 NDNF neurons are crucial for Down state generation ^15^. Consistently, our results show that blocking GABA_B_R-mediated inhibition decreases the coordination of pyramidal neuron activity. However, blocking slow GABA_B_R-mediated inhibition had no impact on pyramidal cell response to spindles nor SWR, while L1 NDNF neuron activity was particularly driven by these oscillations. One explanation for this could be that L1 NDNF-mediated inhibition upon spindles and SWR is mostly driven by the activation of GABA_A_R, whose faster kinetics are more related to the duration of these oscillations ^15^.

### Limitations

Systemic application of a GABA_B_R antagonist would affect other inhibitory synapses in addition to those originating from L1 NDNF cells. Indeed, NDNF interneurons are also present in other brain structures including the hippocampus and the cerebellar cortex ^41^. Likewise, neurogliaform cells are found in other cortical layers such as L2/3, and could participate in the GABA_B_-mediated inhibition^42^. Furthermore, somatostatin-expressing interneurons also activate GABA_B_R in the neocortex ^43^, suggesting that they could participate in the regulation of SWR generation ^44^. Future studies should consider targeting the inhibition of L1 NDNF neurons directly, through an optogenetic or chemogenetic approach in order to clarify the role(s) of L1 NDNF neurons on neuronal coupling.

In summary, our study suggests that L1 NDNF neurons are key regulators of cortical activity during sleep. While recent research has highlighted their role in attention and learning, we propose that they also play a fundamental role in memory consolidation processes. Future behavioral studies will be essential to further clarify their specific contributions to memory consolidation.

## Materials and Methods

### Animals

All experiments were conducted in accordance with United Kingdom Home Office regulations, as outlined in the Animals (Scientific Procedures) Act 1986 Amendment Regulations 2012, and following ethical approval by the University of Cambridge Animal Welfare and Ethical Review Body (AWERB). All animal procedures were performed under Personal and Project licenses held by the authors. Mice were group-housed in conventional open cages with ad libitum access to food and water, maintained on a 12-hour/12-hour light-dark cycle, with temperatures maintained at 22-24 °C and relative humidity kept between 50 % and 55 %. NDNF-Cre mice (Stock number: #028536) were purchased from The Jackson Laboratory and were bred in the animal facility with wild-type C57BL/6J mice purchased from Harlan (Bicester, UK) to keep the transgenic line heterozygous. Although sex differences in the fundamental mechanisms of sleep are not expected, only male mice were used to minimize variance and reduce the number of animals required for statistical analysis. All surgical procedures were carried out in accordance with Home Office standards for aseptic surgery. Mice received analgesic treatment with Meloxicam (2 mg/kg in saline) 30 minutes prior to anesthesia induction using 5 % isoflurane. Once positioned in the stereotaxic frame, the isoflurane concentration was reduced to 1.8 %-2.2 %, and the surgical area was prepared aseptically.

### Viral injections and implant surgeries

The skull was exposed, aligned between bregma and lambda, and holes were drilled above the LFP recording sites (in mm: PFC: anterior-posterior (AP) = 2-2.8, mediolateral (ML) = 0.4; S1: AP = −0.2, ML = 3; CA1: AP = −2.5, ML = 2.5-2.8). Contralaterally to the LFP recording sites, a 3 mm diameter craniotomy was made over the dorsal neocortex, including primary motor, primary somatosensory, parietal, and retrosplenial areas (centered at AP = −1.7, ML = −1.7). Four successive viral injections were performed at depths of 100 µm or 300 µm below the pia to target L1 NDNF neurons (AAV5-Syn-Flex-GCaMP6f-WPRE-SV40) or L2/3 neurons (AAV1-Syn-GCaMP6f-WPRE-SV40), respectively. Eight NDNF-cre mice were used for the main study, including five L1 NDNF mice and three L2/3 mice. The plasmids pAAV.Syn.Flex.GCaMP6f.WPRE.SV40 and pAAV.Syn.GCaMP6f.WPRE.SV40 were a gift from Douglas Kim & the GENIE Project (Addgene plasmid #100833; https://www.addgene.org/100833; RRID and Addgene plasmid #100837; https://www.addgene.org/100837; RRID). A total of 100 nL of virus was delivered at each injection site using a glass pipette, to reduce damage of the dura, attached to a stereotaxic injector. A protective dura gel (Cambridge NeuroTech, UK) was applied over the cranial window. A 3 mm diameter glass coverslip was positioned over the cranial window and secured with glue to the skull. Differential LFP recordings were made using stereotrodes. The stereotrodes consisted of staggered wire electrodes, made of two twisted 125 µm Teflon-coated silver electrodes (AGT0510, World Precision Instruments, Hitchin, UK) with tips spaced 400-600 µm apart. The upper tip of the electrode was implanted in layer 1 and the lower tip in the infragranular layers. For CA1, stereotrodes with tips spaced 200 µm apart were implanted so that the lower tip was inserted 1.2 mm below the pia. S1 and CA1 electrodes were implanted at a 20° angle. The LFP electrodes were fixed to the skull using UV-cured glue (Loctite 303389; Rapid Electronics, UK). Ground and reference silver wires were attached to a stainless steel microscrew placed over the cerebellum. A 125 µm Teflon-coated silver electrode was implanted in the neck muscles to record EMG activity. All wires were connected to a 32-pin Omnetics connector (Genalog, Cranbrook, UK). The connector, electrodes, and an aluminum head bar were secured to the skull using dental cement (Super-Bond C & B; Prestige Dental, Bradford, UK) and dental acrylic cement (Simplex Rapid, Kemdent). Two additional wild-type mice were implanted bilaterally with LFP electrodes only (no viral injections), to test if oscillations propagate simultaneously in both hemisphere (Fig. S4E-J and S5B-D). Following surgery, mice were allowed to recover for a few hours in a heated recovery chamber before being returned to their home cage. Mice received Meloxicam (2 mg/kg) and were weighed daily for five days to ensure proper recovery.

### Miniscope baseplate implantation

Mice were handled daily for three to five consecutive days starting at least five days after the surgery. Using the implanted head bar, mice were progressively habituated to being head-restrained while running on a wheel. Three to four weeks after viral injection, while head-restrained mice were running on the wheel, a recording site with good GCamp6f expression was chosen after scanning the full extent of the cranial-window with a Miniscope V4 ^45^ (https://open-ephys.org/miniscope-v4). Last, the Miniscope baseplate was cemented in isoflurane anesthetized mice as described previously ^46^.

### Recordings in naturally-sleeping mice

Mice (*N* = 5 L1 NDNF mice; *N* = 3 L2/3 mice) were gradually habituated to carry the Miniscope, and to their sleeping cage over 5–7 days. On recording days, to minimize stress, mice were placed in the sleeping cage 2–3 hours before data collection. An RHD 32-channel recording headstage was connected to the Omnetics connector for electrophysiological recordings, which were filtered between 0.1 and 500 Hz and sampled at 2 kHz using the Open Ephys acquisition board and GUI ^47^. Calcium imaging was performed with the Miniscope DAQ v3.3 and Miniscope-DAQ-QT software. For each mouse, LED brightness, focus, and gain were adjusted to optimize neuronal visualization, and these parameters were kept constant over successive recordings. The sample rate ranged from 20 to 30 frames per second and light intensity ranged from 15 to 50 %. For L2/3 neurons, 60-minute-long videos were acquired. For L1 NDNF neurons, 15-minute-long video acquisitions were taken every 30 minutes to minimize photobleaching caused by the higher light intensity needed to image these neurons. To synchronize the videos with the electrophysiological data, a TTL signal was transmitted to the Open Ephys acquisition board via an I/O board for each frame acquired by the Miniscope. Data analysis was conducted offline.

### CGP55,845 injection

The GABA_B_-receptor antagonist, CGP55,845 hydrochloride (Bio-Techne, Abingdon, UK) was injected intraperitoneally at 5 mg/kg (diluted to reach an injection volume of 150-200 µL) 45 min before the beginning of the recordings. As a control, mice were recorded prior to the injection on the same day. In total, 5 mice received the CGP55,845 injection (*N* = 3 L1 NDNF mice; *N* = 2 L2/3 mice).

### Histology

Following the experiment, mice were deeply anesthetized with pentobarbital sodium (90 mg/kg) and, once the absence of reflexes was confirmed, mice were transcardially perfused with 4 % paraformaldehyde in phosphate-buffered saline (PBS). Brains were fixed overnight in 4 % paraformaldehyde and then transferred to 30 % sucrose in PBS solution for 24-48 h. Brains were then embedded, frozen, and cut into 40 µm coronal sections with a cryostat. Sections were stained with G-fluoromount with DAPI prior to mounting and visualization under a confocal microscope (SP8, Leica) using 488-nm Argon laser and analyzed using ImageJ.

### Vigilance state detection

All analyses were performed using custom-made Python scripts. Electrophysiological signals were first down-sampled to 1000 Hz. Signal from LFP electrodes in the superficial layer were subtracted from the signal recorded in the deeper layer in each recording site to remove any distant volume conduction signal from the LFP. Vigilance states were then manually scored on 5-sec epochs, thanks to PFC, S1, and CA1 LPFs and EMG signals. Active wake was characterized by high theta (5-9 Hz) power in CA1 and high EMG amplitude. Quiet wake was identified by moderate EMG amplitude, absence of theta power in CA1, and the presence of slow oscillations (< 4 Hz)—but no sigma power (10-16 Hz)—in S1 and/or PFC. NREM sleep was defined by the presence of both slow oscillations and sigma power in S1 and/or PFC, along with low EMG amplitude. Intermediate sleep was scored when EMG amplitude was low, S1 and/or PFC showed high sigma power, and CA1 exhibited high theta power. REM sleep was characterized by high theta power in CA1, low EMG amplitude, and absence of sigma power in S1 and/or PFC.

### Oscillations detection

The LFP signal from PFC, S1, and CA1was first automatically filtered to remove periods of wakefulness and REM sleep that could alter the detection of oscillations due to potential muscular activity artifacts in the signal. Once done, SWRs were detected in the CA1 LFP, and spindles were extracted from the PFC and S1 signal using the python toolbox Pynapple ^48^. In short, Wavelet decomposition was performed in a 120-200 Hz frequency range for CA1 SWRs and a 10-16 Hz frequency range for PFC and S1 spindles. The mean power for this frequency band was smoothed with a Gaussian kernel of size 0.01 second for SWR and 0.1 second for spindles, and a threshold ranging from 0.05-0.6 was applied depending on the mouse and the recording session. SWR and spindles lasting less than 10 ms and 500 ms, respectively, were removed. S1 and PFC spindles with a minimum overlap of 50 % were identified as S1&PFC spindles. A spindle was considered as coupled with a SWR when at least one SWR started within a maximum of 500 ms before the onset of the spindle or during the spindle.

### Calcium imaging analysis

Videos were processed using MiniAn analysis pipeline ^49^ (https://github.com/denisecailab/minian). In short, raw videos underwent preprocessing, where background vignetting and background fluorescence were corrected, while sensor noise was removed with a median filter. Motion correction was applied using a template-matching algorithm based on cross-correlation between each frame and a reference frame. Local maxima in frame subsets were identified as candidate neurons, and seeds were refined based on signal amplitude and signal-to-noise ratio. These seeds generated initial estimates of the neurons’ spatial footprints and temporal traces. Finally, a constrained nonnegative matrix factorization (CNMF) framework was used to refine the spatial footprints and denoise the temporal traces. Candidate neurons and their transients extracted from MiniAn were manually inspected and non-neuronal shapes were discarded from the analysis. Since mice could be recorded over several days, a cross-registration was performed to identify neurons recorded on multiple videos. Calcium fluorescence intensity was reported in arbitrary units. Activity during vigilance states was evaluated by computing the area under the curve (AUC) for each vigilance state episode. AUC was then normalized by the duration of the vigilance state. AUC during AW, QW, NREM, and REM sleep was used to create a UMAP. Only cells recorded in the four vigilance states under baselined condition (no injected CGP55,845) were included (591/726 L2/3 cells and 217/340 L1 NDNF cells). Parameters used for the UMAP (number of approximate nearest neighbors of 90 and minimum distance between points in low-dimensional space of 0.9) were chosen to preserve the uniformity of the data distribution on the manifold, maintain a locally constant metric, and ensure the manifold’s local connectivity. A Hierarchical Density-Based Spatial Clustering of Applications with Noise (HDBSCAN) (minimum cluster size of 15 and minimum sample in a neighborhood of 1) was then performed on the UMAP for the classification of L1 NDNF cells.

Correlation coefficients (r) between calcium transients were normalized using the z-fisher transform (z) for statistical analysis. Cell assembly was detected as previously described ^50–52^. First, principal component analysis was computed on calcium transients. Assemblies were identified based on principal components whose eigenvalues exceeded the Marčenko-Pastur distribution threshold (Fig. 2D). Subsequently, the fast-independent component analysis (ICA) algorithm was used to extract weight vectors, representing the contribution of each neuron’s activity to each assembly (Fig. 2E). Neurons were classified as members of an assembly if their weight exceeded the assembly’s mean weight by more than 1.5 standard deviations. Cell assemblies with only one cell member were excluded. The activity of each detected cell assembly was computed by averaging the calcium transients of each member. Reactivation events were then identified when the average activity exceeded the mean by 2 standard deviations.

For oscillation analysis, only neurons recorded during at least 10 oscillatory events were considered. Activity upon oscillation onset was computed by subtracting the AUC from 0 to 1 sec after oscillation onset by the AUC from −1 to −0.5 sec before oscillation onset. At the group level, confidence intervals from baselined AUC were used to determine if a group of neurons were responsive to oscillations (confidence intervals excluding 0). At the individual level, neurons were classified as positively or negatively modulated by oscillations when their mean AUC activity from 0 to 1 sec after oscillation onset was superior or inferior to the mean baseline activity (calculated from −1 to −0.5 seconds before onset) by more than 2 standard deviations, respectively. The proportion of modulated cells was compared to a random proportion of unmodulated, positively modulated, and negatively modulated cells computed around randomly chosen time points.

### Statistical Analysis

Statistical analysis was performed using R studio, and the mean +/- the standard error of the mean (SEM) was reported. Two-sample Kolmogorov-Smirnov tests were performed to compare the distribution of two groups. Comparison of sample proportion was performed using the one-proportion Z-test with Yates continuity correction. For multiple group comparisons, one-way ANOVA or two-way ANOVA was performed when one or two factors were included, respectively. Generalized linear mixed-effects model in R studio using the lme4 package and Satterthwaite approximation ^53^ was performed when repeated measurements were included and/or when replicates in one or multiple conditions were missing. Mice, recording sessions, cells, and oscillations were used as random effects when appropriate. Detailed statistics from multiple group comparisons were summarized in Tables S1 and S2. Statistical significance from single group comparison and post-hoc tests were denoted on the graph by asterisks as follows: \**P* < 0.05, \*\**P* < 0.01, \*\*\**P* < 0.001, or detailed in the text.

## Acknowledgments

This work was supported by a Biotechnology and Biological Sciences Research Council (BBSRC) grant (BB/S015922/1) to YAH and OP, an Agence Nationale de la Recherche (ANR) grant (JCJC Solti) to YAH and a Labex Cortex postdoctoral fellowship to AB.

## Supporting Information

**Fig. S1.**
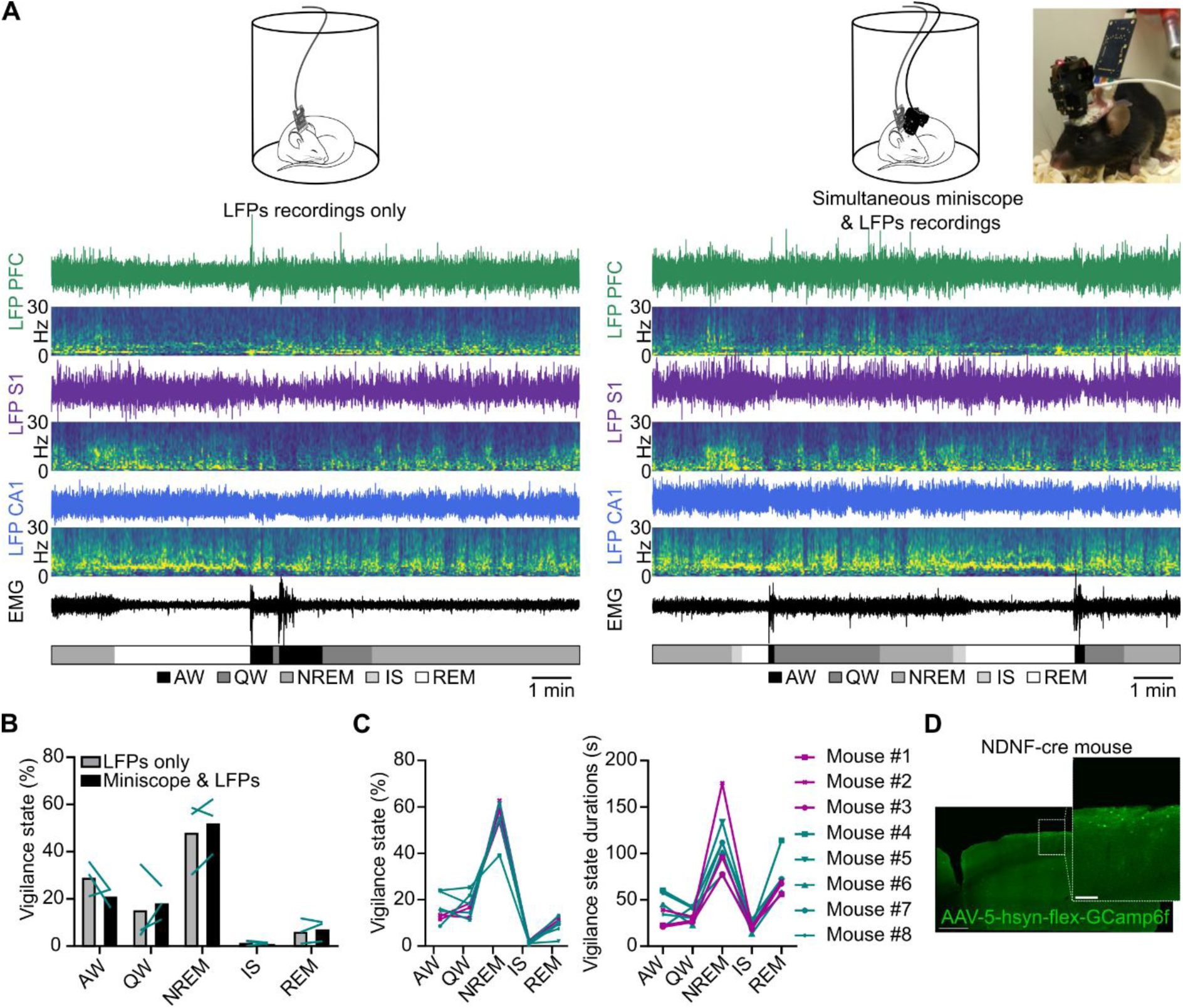
(**A**) Raw LFPs of PFC, S1, CA1, with their respective spectrogram from 0 to 30 Hz, EMG and hypnogram of the same mouse recorded without (left) and with the Miniscope (right). (**B**) Comparison of vigilance states proportion with and without Miniscope recordings. (**C**) Vigilance states proportion (left) and duration (right) per mouse. Pink and blue lines represent L2/3 and L1 NDNF mice, respectively (Proportions: P < 0.01 for all comparisons, except P_AW-QW_, P_AW-REM_, and P_QW-REM_ > 0.05; Durations: P < 0.001 for all comparisons, except P_AW-QW_, P_AW-IS_ and P_IS-QW_ > 0.05). (**D**) Coronal section of a NDNF-Cre mouse brain transduced with a Cre-dependent GCamp6f construct. Scale bars, 100 µm. Generalized linear mixed-effects model (B and C). See Table S2 for statistics. Data are shown as means ± SEM.

**Fig. S2.**
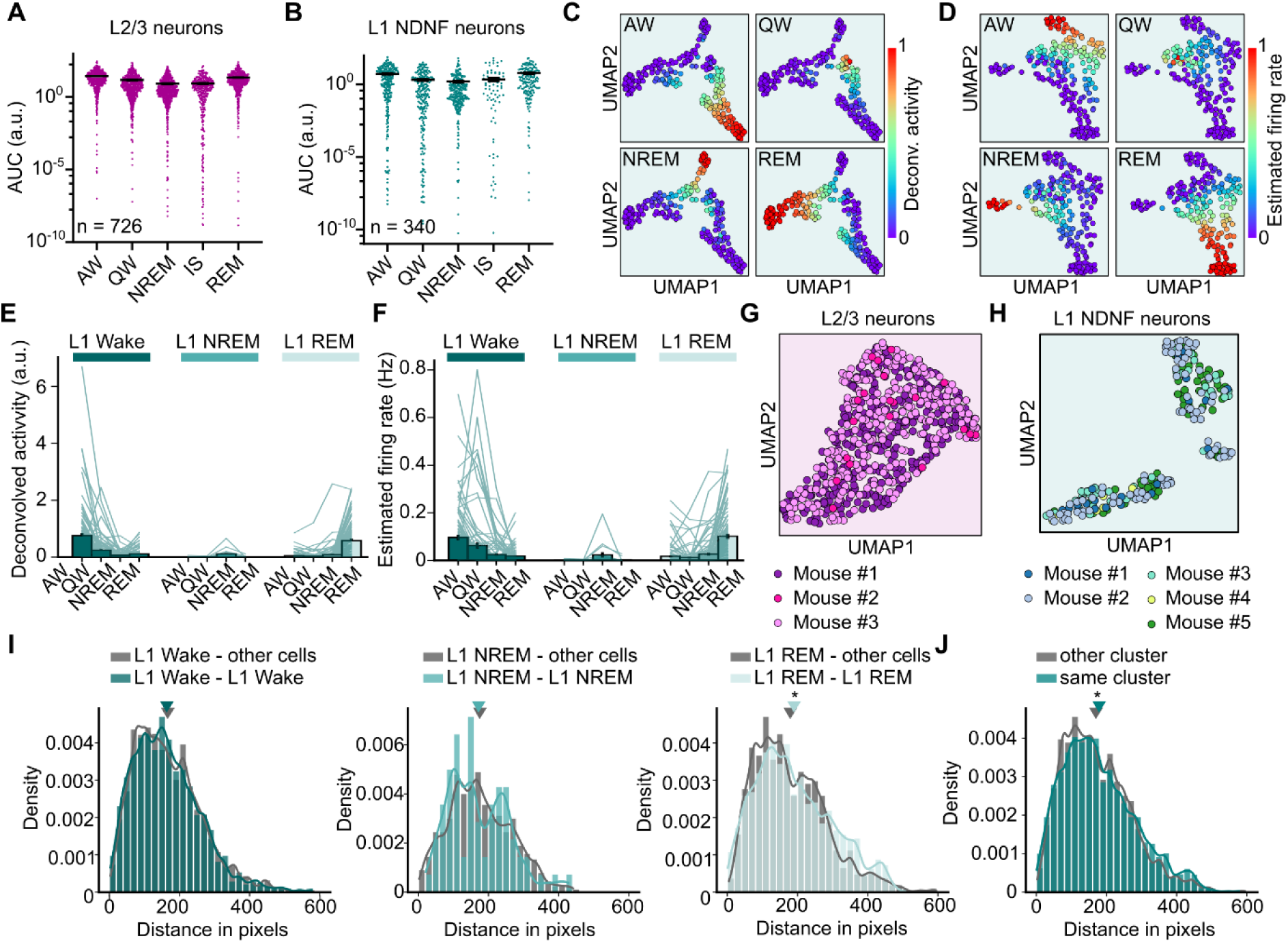
(**A**) Average AUC of the calcium transients of all unique L2/3 neurons according to sleep states (P < 0.05 for all comparisons, except P_QW-REM_, P_QW-IS_ and P_NREM-IS_ > 0.05). (**B**) Same as in A for L1 NDNF neurons (P < 0.05 for all comparisons, except P_AW-REM_, P_IS-NREM_, P_IS-QW_, P_IS-REM_, P_NREM-QW_ > 0.05). (**C**) UMAP based on the deconvolved activity of L1 NDNF cells in each vigilance state. (**D**) UMAP based on the estimated firing rate of L1 NDNF cells in each vigilance state. (**E**) Average deconvolved activity of L1 NDNF neurons according to sleep states and their cluster identity (L1 Wake: P < 0.01 for all comparisons, except P_NREM-REM_ and P_QW-REM_ > 0.05; L1 NREM: P > 0.05 for all comparisons; L1 REM: P < 0.001 for all comparisons, except P_AW-QW,_ P_AW-NREM_ and P_QW-NREM_ > 0.05). (**F**) Average estimated firing rate of L1 NDNF neurons according to sleep states and their cluster identity (L1 Wake: P < 0.01 for all comparisons, except P_NREM-REM_ > 0.05; L1 NREM: P > 0.05 for all comparisons; L1 REM: P < 0.05 for all comparisons, except P_AW-QW_ and P_AW-NREM_ > 0.05). (**G**) UMAP based on the AUC of L2/3 cells per mouse. (**H**) Same as in G for L1 NDNF neurons. (**I**) Distribution of the Euclidian distance between pair of L1 NDNF neurons belonging to the same cluster or not, as a function of their cluster identity. Inverted triangles represent the mean of the distribution. (**J**) Distribution of the Euclidian distance between pair of L1 NDNF neurons belonging to the same cluster or not. Inverted triangles represent the mean of the distribution. *P < 0.05. Generalized linear mixed-effects model (A, B, E, and F) and two-sample Kolmogorov-Smirnov tests (I and J). See Table S2 for statistics. Data are shown as means ± SEM.

**Fig. S3.**
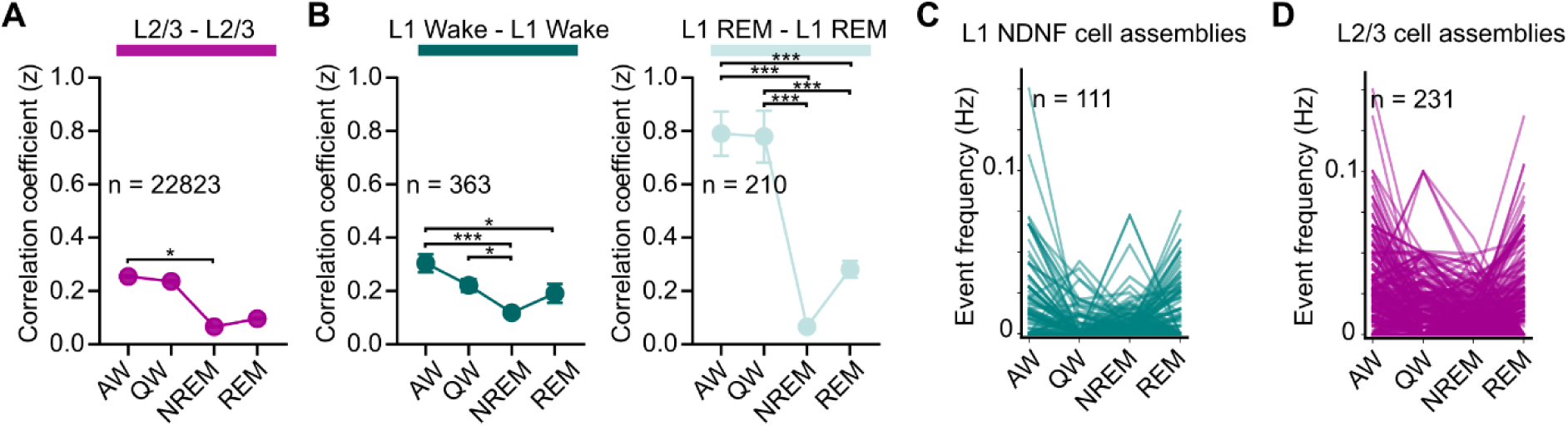
(**A**) Z-transformed correlation coefficient between pair of L2/3 cells recorded in all vigilance states. (**B**) Same as in A for pair of L1 Wake cells (left) and L1 REM cells (right). No L1 NREM cell pairs were recorded in all vigilance sates. (**C**) Mean event frequency of detected cell assemblies of L1 NDNF cells as a function of vigilance states (all comparisons P < 0.05, except P_NREM-QW_ and P_NREM-REM_ > 0.05). (**D**) Same as in C for L2/3 cells assemblies (all comparisons P < 0.05, except P_QW-NREM_ and P_QW-REM_ > 0.05). One-way anova (A, B, C and D). See Table S2 for statistics. Data are shown as means ± SEM.

**Fig. S4.**
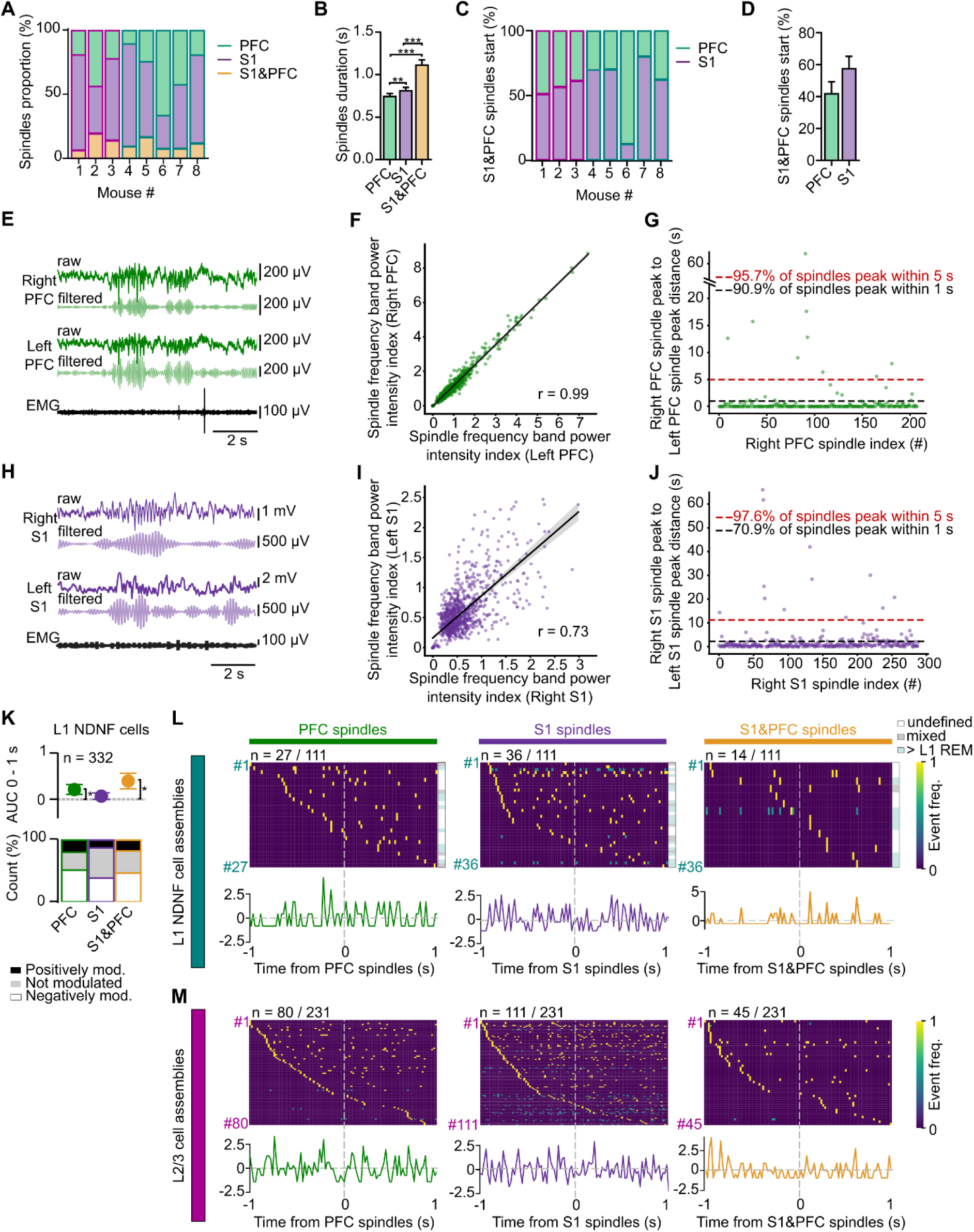
(**A**) Average proportion of spindles detected depending on their locations per mouse. Pink and blue outlines represent L2/3 and L1 NDNF mice, respectively. (**B**) Spindle durations depending on their locations (*N* = 8 mice). (**C**) Proportion of S1&PFC spindles starting first in S1 or PFC per mouse. Pink and blue outlines represent L2/3 and L1 NDNF mice, respectively. (**D**) Proportion of S1&PFC spindles starting first in S1 or PFC (*N* = 8 mice). (**E**) Example of right and left, raw and filtered (10-16 Hz) PFC LFPs recorded simultaneously during NREM sleep. (**F**) Pearson correlation on randomly extracted 10000 data points from right and left averaged power intensity of PFC LFPs on a mouse. (**G**) Computation of the minimal distance between right and left PFC spindles (N = 1 mouse). (**H**-**J**) Same as E to G for S1 LFP. (**K**) Average baselined AUC (a.u.) of all L1 NDNF cells after spindle onset (top). Proportion of unmodulated, positively modulated and negatively modulated cells by spindles as a function of spindles location (bottom). (**L**) Normalized event frequency of L1 NDNF cell assemblies one second before and after spindle onset. Only active cell assembly are represented on the heatmap (top). On the right of the heatmaps, a color code indicates the nature of the cell assembly (undefined, mixed or predominately composed of L1 REM cells). Average event frequency around oscillations (bottom). No significant activity was detected (P > 0.05). (**M**) Same as in L for L2/3 cell assemblies. No significant activity was detected (P > 0.05). Generalized linear mixed-effects model (B, D and K) and permutations tests (L and M). See Table S2 for statistics. Data are shown as means ± SEM.

**Fig. S5.**
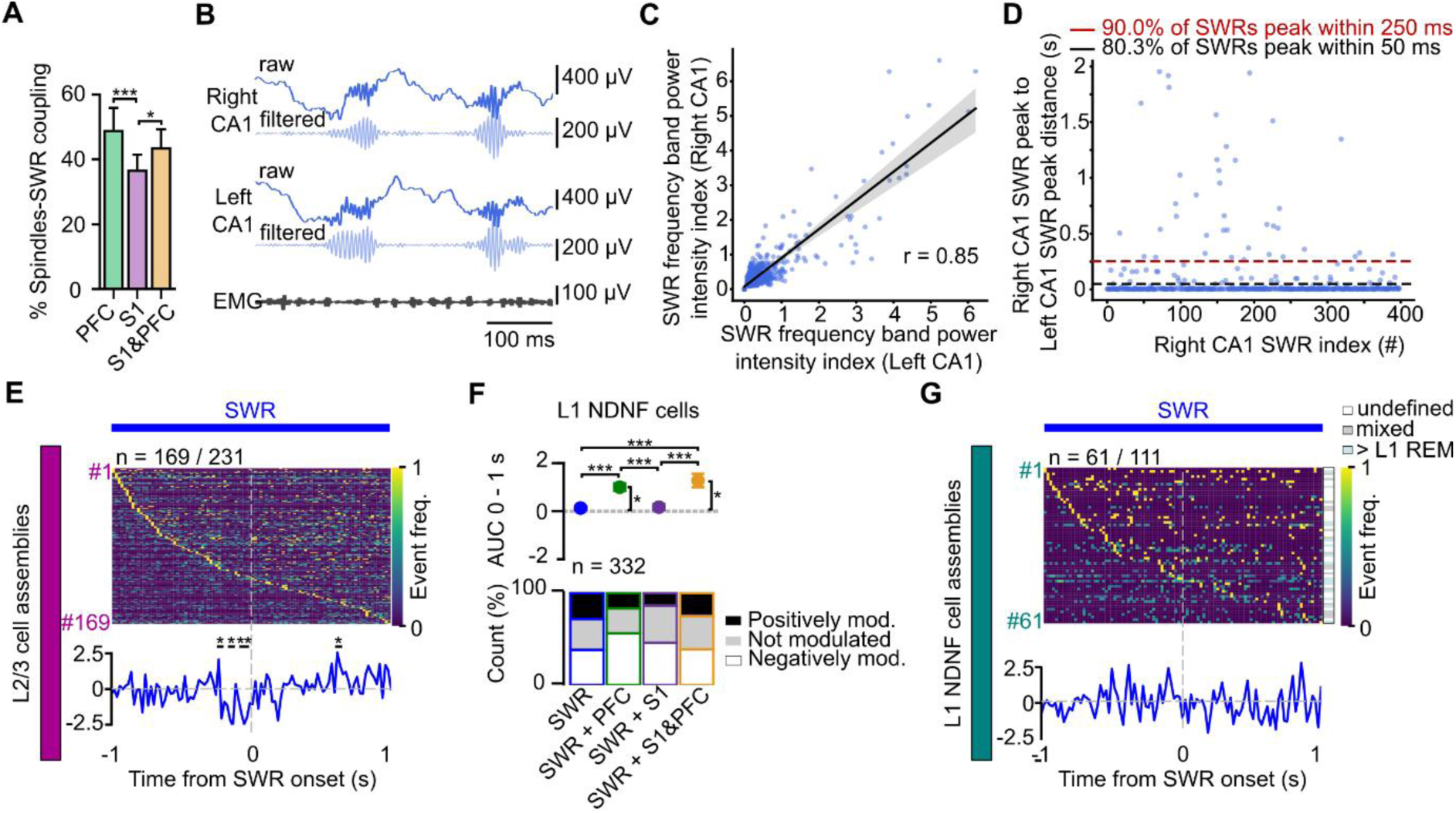
(**A**) Percentage of spindles coupled with SWRs depending on the spindle locations (*N* = 8 mice). (**B**) Example of right and left, raw and filtered (120-200 Hz) CA1 LFPs recorded simultaneously during NREM sleep. (**C**) Pearson correlation on randomly extracted 10000 data points from right and left averaged power intensity of CA1 LFPs on a mouse. (**D**) Computation of the minimal distance between right and left CA1 SWR (N = 1 mouse). (**E**) Normalized event frequency of L2/3 cell assemblies one second before and after SWR onset. Only active cell assembly are represented on the heatmap (top). Average event frequency around oscillations (bottom). (**F**) Average baselined AUC (a.u.) of all L1 NDNF neurons after SWR onset (top). Proportion of unmodulated, positively modulated and negatively modulated cells by SWR as a function of its coupling (bottom). (**G**) Same as in E for L1NDNF cell assemblies. On the right of the heatmaps, a color code indicates the nature of the cell assembly (undefined, mixed or predominately composed of L1 REM cells). *P < 0.05, ***P < 0.001. Generalized linear mixed-effects model (A and F) and permutations tests (E and G). See Table S2 for statistics. Data are shown as means ± SEM.

**Fig. S6.**
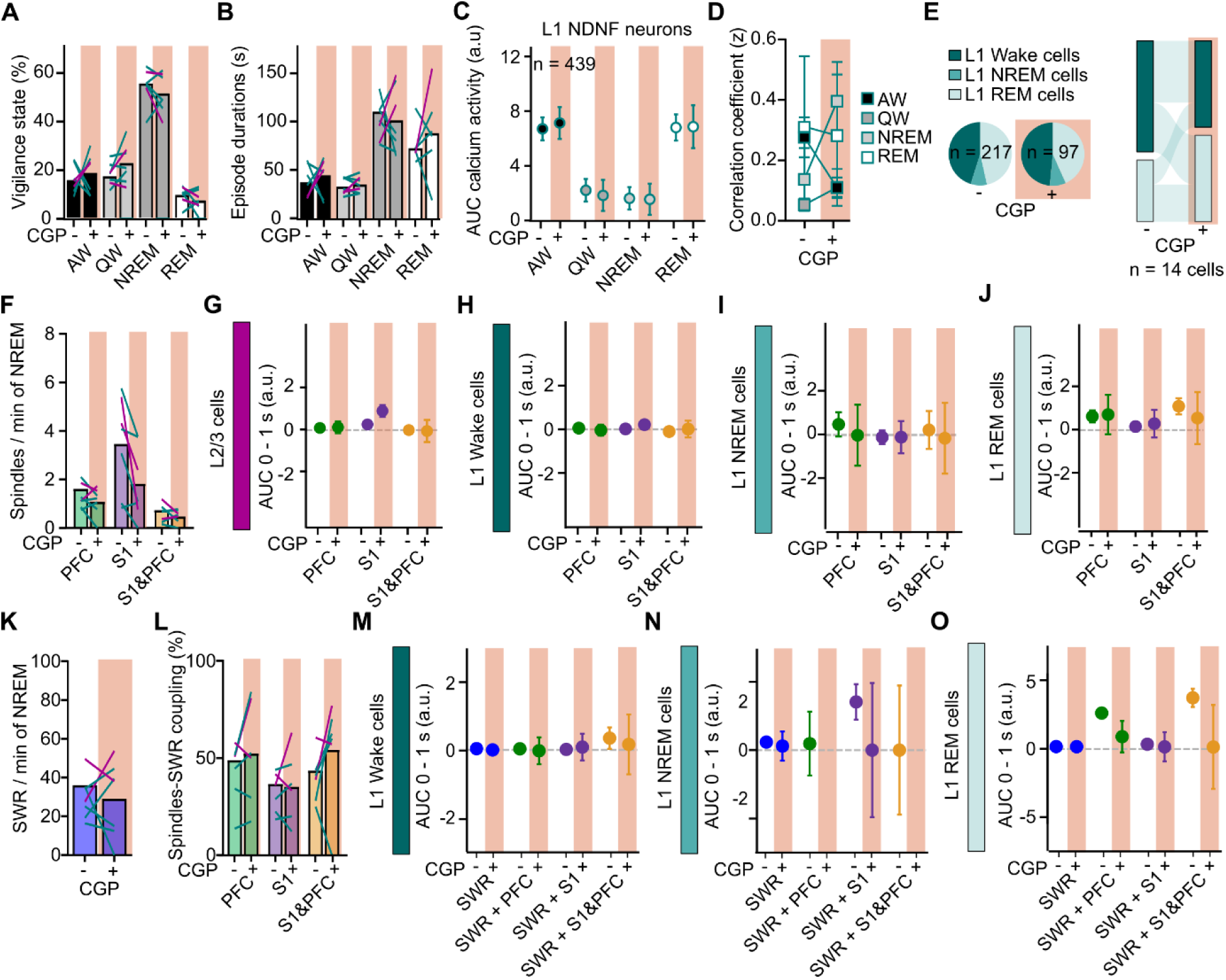
Impact of CGP55,845 injection on the proportion of vigilance state (**A**) and the episode durations (**B**) during the recording sessions. Each lines represents a mouse before and after CGP55,845 injection. Pink and blue lines represent L2/3 and L1 NDNF mice, respectively. (**C**) Average AUC of L1 NDNF neuron calcium activity before and after CGP55,845 injection according to vigilance states. (**D**) Evolution of the correlation coefficient (z) before and after CGP55,845 injection per vigilance states between L1 NDNF cells. (**E**) Proportion of the three subpopulations of L1 NDNF neurons under baseline condition versus after CGP55,845 injection (left). Classification of L1 NDNF neurons recorded in all vigilance states before and after CGP55,845 injection (right). (**F**) Spindle rate during NREM sleep before and after CGP55,845 injection. (**G**) Mean baselined activity of L2/3 neurons 1 second after spindle onset depending on the spindle locations under baseline condition versus after CGP55,845 injection. (**H**) Same as in G for L1 Wake cells. (**I**) Same as in G for L1 NREM cells. (**J**) Same as in G for L1 REM cells. (**K**) SWR rate during NREM sleep before and after CGP55,845 injection. (**L**) Percentage of spindles coupled with SWRs depending on the spindle locations before and after CGP55,845 injection. (**M**) Mean baselined activity of L1 Wake neurons 1 second after SWR onset depending on the SWR coupling under baseline condition versus after CGP55,845 injection. (**N**) Same as in M for L1 NREM cells. (**O**) Same as in M for L1 REM cells. Generalized linear mixed-effects model (A, B, C, F-O) and two-way anova (D). See Table S2 for statistics. Data are shown as means ± SEM.

**Table S1.** Statistics related to main figures.

| Fig | Tested variable | Method | Statistics value | Degrees of freedom | P values | Post-hoc tests |
| --- | --- | --- | --- | --- | --- | --- |
| Fig. 1G | AUC of L2/3 cells (a.u) | Linear Mix Model (fix: Substates, random: Mouse ID, session ID, substate ID, Unit ID) | 30.8 | 3, 422 | < 2.2e-16 | 6 x Tukey |
| Fig. 1L | AUC of L1 NDNF cells (a.u) | Linear Mix Model (fix: Substates, random: Mouse ID, session ID, substate ID, Unit ID) | 23.1 | 3, 364 | 1.04e-13 | 6 x Tukey |
| Fig. 1O | AUC of L1 Wake, NREM and REM cells (a.u) | Linear Mix Model (fix: Substates*Cluster ID, random: Mouse ID, session ID, Unit ID) | Substates: 9.9<br>Cluster ID: 3.1<br>Interaction: 33.4 | Substates: 3, 673<br>Cluster ID: 2, 308<br>Interaction: 6, 4956 | Substates: 2.04e-06<br>Cluster ID: 0.045<br>Interaction: < 2.2e-16 | 18 x Tukey |
| Fig. 2C | Correlation coefficient of L1 NDNF cells (z) | Two-way ANOVA | Pairs: 41.6<br>Cluster ID: 2.45<br>Interaction: 3.33 | Pairs: 1, 4320<br>Cluster ID: 2, 4319<br>Interaction: 2, 4319 | Pairs: 1.24e-10<br>Cluster ID: 0.086<br>Interaction: 0.036 | 3 x Tukey |
| Fig. 3D | AUC of L2/3 cells after spindles onset (a.u.) | Linear Mix Model (fix: Spindles location ID, random: Mouse ID, session ID, Unit ID, Spindles ID) | 1.4 | 2, 1074 | 0.24 | - |
| Fig. 3E | AUC of L1 Wake cells after spindles onset (a.u.) | Linear Mix Model (fix: Spindles location, random: Mouse ID, session ID, Unit ID, Spindles ID) | 0.66 | 2, 592 | 0.51 | - |
| Fig. 3F | AUC of L1 NREM cells after spindles onset (a.u.) | Linear Mix Model (fix: Spindles location, random: Mouse ID, session ID, Unit ID, Spindles ID) | 0.35 | 2, 313 | 0.70 | - |
| Fig. 3G | AUC of L1 REM cells after spindles onset (a.u.) | Linear Mix Model (fix: Spindles location, random: Mouse ID, session ID, Unit ID, Spindles ID) | 4.2 | 2, 262 | 0.017 | 3 x Tukey |
| Fig. 4B | AUC of L2/3 cells after SWR onset (a.u.) | Linear Mix Model (fix: SWR coupling, random: Mouse ID, session ID, Unit ID, SWR ID) | 0.55 | 3, 5604 | 0.65 | - |
| Fig. 4C | AUC of L1 Wake cells after SWR onset (a.u.) | Linear Mix Model (fix: SWR coupling, random: Mouse ID, session ID, Unit ID, SWR ID) | 0.32 | 3, 5274 | 0.81 | - |
| Fig. 4D | AUC of L1 NREM cells after SWR onset (a.u.) | Linear Mix Model (fix: SWR coupling, random: Mouse ID, session ID, Unit ID, SWR ID) | 1.6 | 3, 1970 | 0.18 | - |
| Fig. 4E | AUC of L1 REM cells after SWR onset (a.u.) | Linear Mix Model (fix: SWR coupling, random: Mouse ID, session ID, Unit ID, SWR ID) | 30.4 | 3, 26892 | < 2.2e-16 | 6 x Tukey |
| Fig. 5C | AUC of L2/3 cells | Linear Mix Model (fix: Substates*Drug, random: Mouse ID, session ID, substate ID, Unit ID) | Substates: 13.1<br>Drug: 0.2<br>Interaction: 0.12 | Substates: 3, 299<br>Drug: 1, 19<br>Interaction: 3, 299 | Substates: 4.5e-8<br>Drug: 0.63<br>Interaction: 0.94 | - |
| Fig. 5D | Correlation coefficient of L2/3 cells (z) | Two-way ANOVA | Substates: 17.0<br>Drug: 120.4<br>Interaction: 11.5 | Substates: 3, 13358<br>Drug: 1, 13360<br>Interaction: 3, 13358 | Substates: 5.29e-11<br>Drug: 6.82e-28<br>Interaction: 1.7e-7 | - |
| Fig. 5E | AUC of L2/3 cells after spindles onset (a.u.) | Linear Mix Model (fix: SWR coupling *Drug, random: Mouse ID, session ID, Unit ID, SWR ID) | Coupling: 0.18<br>Drug: 0.0089<br>Interaction: 0.36 | Coupling: 3, 9321<br>Drug: 1, 14<br>Interaction: 3, 9319 | Coupling: 0.91<br>Drug: 0.92<br>Interaction: 0.78 | - |

**Table S2.** Statistics related to supplementary figures.

| Fig | Tested variable | Method | Statistics value | df | P values | Post-hoc tests |
| --- | --- | --- | --- | --- | --- | --- |
| Fig. S1B | Vigilance state (%) | Linear Mix Model<br>(fix: Substates<br>*Miniscope, random:<br>Mouse ID) | Substates:<br>16.0<br>Miniscope:<br>1.0<br>Interaction:<br>0.32 | Substates:<br>1, 2<br>Miniscope:<br>1, 2<br>Interaction:<br>1, 2 | Substates:<br>0.048<br>Miniscope:<br>0.97<br>Interaction:<br>0.39 | - |
| Fig. S1C (left) | Vigilance state (%) | Linear Mix Model<br>(fix: Substates,<br>random: Mouse ID, session ID) | 166.3 | 4, 176 | < 2.2e-16 | 6 x Tukey |
| Fig. S1C (right) | Vigilance duration (s) | Linear Mix Model<br>(fix: Substates,<br>random: Mouse ID, session ID) | 89.6 | 4, 1161 | < 2.2e-16 | 6 x Tukey |
| Fig. S2A | AUC of L2/3 cells (a.u) | Linear Mix Model<br>(fix: Substates,<br>random: Mouse ID, session ID, Substate ID, Unit ID) | 24.8 | 4, 452 | < 2.2e-16 | 6 x Tukey |
| Fig. S2B | AUC of L1 NDNF cells (a.u) | Linear Mix Model<br>(fix: Substates,<br>random: Mouse ID, session ID, substate ID, Unit ID) | 19.4 | 4, 416 | 1.3e-11 | 6 x Tukey |
| Fig. S2E | Deconvolved activity of L1 Wake, NREM, REM cells (a.u) | Linear Mix Model<br>(fix: Substates*Cluster ID, random: Mouse ID, session ID, substate ID, Unit ID) | Substates:<br>3.4<br>Cluster ID:<br>6.1<br>Interaction:<br>26.6 | Substates:<br>3, 935<br>Cluster ID:<br>2, 376<br>Interaction:<br>6, 5038 | Substates:<br>0.017<br>Cluster ID:<br>0.002<br>Interaction:<br>< 2.2e-16 | 18 x Tukey |
| Fig. S2F | Estimated firing rate of L1 Wake, NREM, REM cells (a.u) | Linear Mix Model<br>(fix: Substates*Cluster ID, random: Mouse ID, session ID, substate ID, Unit ID) | Substates:<br>5.1<br>Cluster ID:<br>5.7<br>Interaction:<br>43.6 | Substates:<br>3, 783<br>Cluster ID:<br>2, 214<br>Interaction:<br>6, 4919 | Substates:<br>0.0018<br>Cluster ID:<br>0.003<br>Interaction:<br>< 2.2e-16 | 18 x Tukey |
| Fig. S3A | Correlation coefficient of L2/3 cells (z) | One-way ANOVA | 83.0 | 3, 105 | 1.5e-27 | 6 x Tukey |
| Fig. S3B (left) | Correlation coefficient of L1 Wake cells (z) | One-way ANOVA | 10.2 | 3, 688 | 1.4e-6 | 6 x Tukey |
| Fig. S3B (right) | Correlation coefficient of L1 REM cells (z) | One-way ANOVA | 50.7 | 3, 321 | 7.62e-27 | 6 x Tukey |
| Fig. S3C | Event frequency L1 NDNF cell assemblies (Hz) | One-way ANOVA | 15.96 | 3, 616 | 5.26e-10 | 6 x Tukey |
| Fig. S3D | Event frequency L2/3 cell assemblies (Hz) | One-way ANOVA | 19.55 | 3, 924 | 2.71e-12 | 6 x Tukey |
| Fig. S4B | Spindles durations (s) | Linear Mix Model (fix: Spindles location, random: Mouse ID, session ID) | 202 | 2, 3792 | < 2.2e-16 | 3 x Tukey |
| Fig. S4D | S1&PFC spindles start (%) | Linear Mix Model (fix: Start Location, random: Mouse ID, session ID) | 3.0 | 1,80 | 0.08 | - |
| Fig. S4K | AUC of L1 NDNF cells after spindles onset (a.u.) | Linear Mix Model (fix: Spindles location, random: Mouse ID, session ID, Unit ID, Spindles ID) | 2.1 | 2, 713 | 0.13 | - |
| Fig. S5A | Spindles-SWR coupling (%) | Linear Mix Model (fix: Spindles location, random: Mouse ID, session ID) | 9.0 | 2, 76 | 0.00030 | 3 x Tukey |
| Fig. S5F | AUC of L1 NDNF cells after SWR onset (a.u.) | Linear Mix Model (fix: SWR coupling, random: Mouse ID, session ID, Unit ID, SWR ID) | 18.5 | 3, 7462 | 6.38e-12 | 6 x Tukey |
| Fig. S6A | Vigilance states proportion (%) | Linear Mix Model (fix: Substates*Drug, random: Mouse ID, session ID) | Substates: 114.3<br>Drug: 0.0002<br>Interaction: 0.7 | Substates: 3, 182<br>Drug: 1, 182<br>Interaction: 3, 182 | Substates: < 2.2e-16<br>Drug: 0.98<br>Interaction: 0.54 | - |
| Fig. S6B | Vigilance states durations (s) | Linear Mix Model (fix: Substates*Drug, random: Mouse ID, session ID) | Substates: 75.5<br>Drug: 0.2<br>Interaction: 0.7 | Substates: 3, 1382<br>Drug: 1, 57<br>Interaction: 3, 1382 | Substates: < 2.2e-16<br>Drug: 0.62<br>Interaction: 0.55 | - |
| Fig. S6C | AUC of L1 NDNF cells | Linear Mix Model (fix: Substates*Drug, random: Unit ID) | Substates: 69.8<br>Drug: 0.0001<br>Interaction: 0.3 | Substates: 3, 8017<br>Drug: 1, 26<br>Interaction: 3, 8019 | Substates: < 2.2e-16<br>Drug: 0.99<br>Interaction: 0.82 | - |
| Fig. S6D | Correlation coefficient of L1 NDNF cells (z) | Two-way ANOVA | Substates: 2.4<br>Drug: 0.1<br>Interaction: 2.8 | Substates: 3<br>Drug: 1<br>Interaction: 3 | Substates: 0.064<br>Drug: 0.71<br>Interaction: 0.04 | 4 x Tukey |
| Fig. S6F | Spindles rate during NREM | Linear Mix Model (fix: Drug*Spindle location, random: Mouse ID) | Drug: 2.3<br>Spindles location: 20.7<br>Interaction: 0.7 | Drug: 1, 137<br>Spindles location: 2, 131<br>Interaction: 2, 131 | Drug: 0.13<br>Spindles location: 1.54e-8<br>Interaction: 0.48 | - |
| Fig. S6G | AUC of L2/3 cells after spindle onset (a.u.) | Linear Mix Model (fix: Drug*Spindle location, random: Mouse ID, session ID, Unit ID, Spindle ID) | Drug: 0.8<br>Spindles location: 3.4<br>Interaction: 1.3 | Drug: 1, 1077<br>Spindles location: 2, 1048<br>Interaction: 2, 1048 | Drug: 0.36<br>Spindles location: 0.032<br>Interaction: 0.27 | - |
| Fig. S6H | AUC of L1 Wake cells after spindle onset (a.u.) | Linear Mix Model (fix: Drug*Spindle location, random: Mouse ID, session ID, Unit ID, Spindle ID) | Drug: 0.2<br>Spindles location: 0.7<br>Interaction: 0.6 | Drug: 1, 2407<br>Spindles location: 2, 5213<br>Interaction: 2, 6016 | Drug: 0.68<br>Spindles location: 0.66<br>Interaction: 0.63 | - |
| Fig. S6I | AUC of L1 NREM cells after spindle onset (a.u.) | Linear Mix Model (fix: Drug*Spindle location, random: Mouse ID, session ID, Unit ID, Spindle ID) | Drug: 0.1<br>Spindles location: 0.08<br>Interaction: 0.05 | Drug: 1, 493<br>Spindles location: 2, 501<br>Interaction: 2, 501 | Drug: 0.72<br>Spindles location: 0.92<br>Interaction: 0.94 | - |
| Fig. S6J | AUC of L1 REM cells after spindle onset (a.u.) | Linear Mix Model (fix: Drug*Spindle location, random: Mouse ID, session ID, Unit ID, Spindle ID) | Drug: 0.04<br>Spindles location: 0.6<br>Interaction: 0.1 | Drug: 1, 1274<br>Spindles location: 2, 4889<br>Interaction: 2, 5600 | Drug: 0.56<br>Spindles location: 0.85<br>Interaction: 0.87 | - |
| Fig. S6K | SWR rate during NREM | Linear Mix Model (fix: Drug, random: Mouse ID) | 1.4 | 1, 40 | 0.24 | - |
| Fig. S6L | Spindles-SWR coupling (%) | Linear Mix Model (fix: Drug*Spindles location, random: Mouse ID, session ID) | Drug: 2.3<br>Spindles location: 11.5<br>Interaction: 1.9 | Drug: 1, 41<br>Spindles location: 2, 88<br>Interaction: 2, 88 | Drug: 0.13<br>Spindles location: 3.54e-5<br>Interaction: 0.15 | - |
| Fig. S6M | AUC of L1 Wake cells after SWR onset (a.u.) | Linear Mix Model (fix: Drug*SWR coupling, random: Mouse ID, session ID, Unit ID, SWR ID) | Drug: 0.03<br>Coupling: 0.09<br>Interaction: 0.03 | Drug: 1, 18982<br>Coupling: 3, 27717<br>Interaction: 3, 28239 | Drug: 0.85<br>Coupling: 0.96<br>Interaction: 0.99 | - |
| Fig. S6N | AUC of L1 NREM cells after SWR onset (a.u.) | Linear Mix Model (fix: Drug* SWR coupling, random: Mouse ID, session ID, Unit ID, SWR ID) | Drug: 0.53<br>Coupling: 0.26<br>Interaction: 0.38 | Drug: 1, 4114<br>Coupling: 3, 4124<br>Interaction: 1, 4114 | Drug: 0.46<br>Coupling: 0.85<br>Interaction: 0.53 | - |
| Fig. S6O | AUC of L1 REM after SWR onset (a.u.) | Linear Mix Model (fix: Drug* SWR coupling, random: Mouse ID, session ID, Unit ID, SWR ID) | Drug: 2.2<br>Coupling: 2.9<br>Interaction: 1.2 | Drug: 1, 1513<br>Coupling: 3, 28456<br>Interaction: 3, 28322 | Drug: 0.13<br>Coupling: 0.03<br>Interaction: 0.32 | - |

## References

1. Born, J. & Wilhelm, I. System consolidation of memory during sleep. Psychological Research 76, 192–203 (2012).

2. Stickgold, R. & Walker, M. Memory consolidation and reconsolidation: what is the role of sleep? Trends in Neurosciences 28, 408–415 (2005).

3. Sirota, A., Csicsvari, J., Buhl, D. & Buzsáki, G. Communication between neocortex and hippocampus during sleep in rodents. Proc. Natl. Acad. Sci. U.S.A. 100, 2065–2069 (2003).

4. Battaglia, F. P., Sutherland, G. R. & McNaughton, B. L. Hippocampal sharp wave bursts coincide with neocortical “up-state” transitions. Learn. Mem. 11, 697–704 (2004).

5. Ngo, H.-V., Fell, J. & Staresina, B. Sleep spindles mediate hippocampal-neocortical coupling during long-duration ripples. eLife 9, e57011 (2020).

6. Pedrosa, R. et al. Hippocampal ripples coincide with “up-state” and spindles in retrosplenial cortex. Cerebral Cortex 34, bhae083 (2024).

7. Schreiner, T., Petzka, M., Staudigl, T. & Staresina, B. P. Endogenous memory reactivation during sleep in humans is clocked by slow oscillation-spindle complexes. Nat Commun 12, 3112 (2021).

8. Siapas, A. G. & Wilson, M. A. Coordinated Interactions between Hippocampal Ripples and Cortical Spindles during Slow-Wave Sleep. Neuron 21, 1123–1128 (1998).

9. Staresina, B. P., Niediek, J., Borger, V., Surges, R. & Mormann, F. How coupled slow oscillations, spindles and ripples coordinate neuronal processing and communication during human sleep. Nat Neurosci 26, 1429–1437 (2023).

10. Maingret, N., Girardeau, G., Todorova, R., Goutierre, M. & Zugaro, M. Hippocampo-cortical coupling mediates memory consolidation during sleep. Nat Neurosci 19, 959–964 (2016).

11. Pardi, M. B., Schroeder, A. & Letzkus, J. J. Probing top-down information in neocortical layer 1. Trends in Neurosciences 46, 20–31 (2023).

12. Schuman, B., Dellal, S., Prönneke, A., Machold, R. & Rudy, B. Neocortical Layer 1: An Elegant Solution to Top-Down and Bottom-Up Integration. Annu. Rev. Neurosci. 44, 221–252 (2021).

13. Hartung, J., Schroeder, A., Péréz Vázquez, R. A., Poorthuis, R. B. & Letzkus, J. J. Layer 1 NDNF interneurons are specialized top-down master regulators of cortical circuits. Cell Reports 43, 114212 (2024).

14. Schuman, B. et al. Four Unique Interneuron Populations Reside in Neocortical Layer 1. J. Neurosci. 39, 125–139 (2019).

15. Hay, Y. A. et al. Thalamus mediates neocortical Down state transition via GABAB-receptor-targeting interneurons. Neuron 109, 2682–2690.e5 (2021).

16. Tamás, G., Simon, A. L., Anna & Szabadics, J. Identified Sources and Targets of Slow Inhibition in the Neocortex. Science 299, 1902–1905 (2003).

17. Jiang, X., Wang, G., Lee, A. J., Stornetta, R. L. & Zhu, J. J. The organization of two new cortical interneuronal circuits. Nat Neurosci 16, 210–218 (2013).

18. Schulz, J. M., Kay, J. W., Bischofberger, J. & Larkum, M. E. GABAB Receptor-Mediated Regulation of Dendro-Somatic Synergy in Layer 5 Pyramidal Neurons. Front. Cell. Neurosci. 15, 718413 (2021).

19. Cohen-Kashi Malina, K., et al. NDNF interneurons in layer 1 gain-modulate whole cortical columns according to an animal’s behavioral state. Neuron 109, 2150–2164.e5 (2021).

20. Abs, E. et al. Learning-Related Plasticity in Dendrite-Targeting Layer 1 Interneurons. Neuron 100, 684–699.e6 (2018).

21. Pardi, M. B. et al. A thalamocortical top-down circuit for associative memory. Science 370, 844–848 (2020).

22. Li, B. et al. Circuit mechanism for suppression of frontal cortical ignition during NREM sleep. Cell 186, 5739–5750.e17 (2023).

23. Geva-Sagiv, M. et al. Augmenting hippocampal–prefrontal neuronal synchrony during sleep enhances memory consolidation in humans. Nat Neurosci 26, 1100–1110 (2023).

24. Helfrich, R. F. et al. Bidirectional prefrontal-hippocampal dynamics organize information transfer during sleep in humans. Nat Commun 10, 3572 (2019).

25. Chambers, A. R., Berge, C. N. & Vervaeke, K. Cell-type-specific silence in thalamocortical circuits precedes hippocampal sharp-wave ripples. Cell Rep 40, 111132 (2022).

26. Kohl, M. M. & Paulsen, O. The Roles of GABAB Receptors in Cortical Network Activity. In Advances in Pharmacology vol. 58 205–229 (Elsevier, 2010).

27. Niethard, N. et al. Sleep-Stage-Specific Regulation of Cortical Excitation and Inhibition. Current Biology 26, 2739–2749 (2016).

28. Brécier, A., Borel, M., Urbain, N. & Gentet, L. J. Vigilance and Behavioral State-Dependent Modulation of Cortical Neuronal Activity throughout the Sleep/Wake Cycle. J. Neurosci. 42, 4852– 4866 (2022).

29. Vazquez, J. & Baghdoyan, H. A. Basal forebrain acetylcholine release during REM sleep is significantly greater than during waking. American Journal of Physiology-Regulatory, Integrative and Comparative Physiology 280, R598–R601 (2001).

30. Poorthuis, R. B. et al. Rapid Neuromodulation of Layer 1 Interneurons in Human Neocortex. Cell Reports 23, 951–958 (2018).

31. Christophe, E. et al. Two Types of Nicotinic Receptors Mediate an Excitation of Neocortical Layer I Interneurons. Journal of Neurophysiology 88, 1318–1327 (2002).

32. Hay, Y. A., Lambolez, B. & Tricoire, L. Nicotinic Transmission onto Layer 6 Cortical Neurons Relies on Synaptic Activation of Non-α7 Receptors. Cereb. Cortex 26, 2549–2562 (2016).

33. Hay, Y. A., Naudé, J., Faure, P. & Lambolez, B. Target Interneuron Preference in Thalamocortical Pathways Determines the Temporal Structure of Cortical Responses. Cereb Cortex 29, 2815–2831 (2019).

34. Bugeon, S. et al. A transcriptomic axis predicts state modulation of cortical interneurons. Nature 607, 330–338 (2022).

35. Anderer, P. et al. Low-resolution brain electromagnetic tomography revealed simultaneously active frontal and parietal sleep spindle sources in the human cortex. Neuroscience 103, 581–592 (2001).

36. Piantoni, G., Halgren, E. & Cash, S. S. Spatiotemporal characteristics of sleep spindles depend on cortical location. NeuroImage 146, 236–245 (2017).

37. Kim, D., Hwang, E., Lee, M., Sung, H. & Choi, J. H. Characterization of Topographically Specific Sleep Spindles in Mice. Sleep 38, 85–96 (2015).

38. Bonjean, M. et al. Corticothalamic Feedback Controls Sleep Spindle Duration In Vivo. Journal of Neuroscience 31, 9124–9134 (2011).

39. Blanco-Duque, C. et al. Oscillatory-Quality of sleep spindles links brain state with sleep regulation and function. Sci. Adv. 10, eadn6247 (2024).

40. Opalka, A. N., Huang, W.-Q., Liu, J., Liang, H. & Wang, D. V. Hippocampal Ripple Coordinates Retrosplenial Inhibitory Neurons during Slow-Wave Sleep. Cell Rep 30, 432–441.e3 (2020).

41. Kuang, X.-L. et al. Spatio-temporal expression of a novel neuron-derived neurotrophic factor (NDNF) in mouse brains during development. BMC Neurosci 11, 137 (2010).

42. Kawaguchi, Y. Physiological subgroups of nonpyramidal cells with specific morphological characteristics in layer II/III of rat frontal cortex. J. Neurosci. 15, 2638–2655 (1995).

43. Kanigowski, D., Bogaj, K., Barth, A. L. & Urban-Ciecko, J. Somatostatin-expressing interneurons modulate neocortical network through GABAb receptors in a synapse-specific manner. Sci Rep 13, 8780 (2023).

44. Zhang, C.-L., Sontag, L., Gómez-Ocádiz, R. & Schmidt-Hieber, C. Learning-dependent gating of hippocampal inputs by frontal interneurons. Proc. Natl. Acad. Sci. U.S.A. 121, e2403325121 (2024).

45. Ghosh, K. K. et al. Miniaturized integration of a fluorescence microscope. Nat Methods 8, 871–878 (2011).

46. Jarzebowski, P., Hay, Y. A., Grewe, B. F. & Paulsen, O. Different encoding of reward location in dorsal and intermediate hippocampus. Current Biology 32, 834–841.e5 (2022).

47. Siegle, J. H. et al. Open Ephys: an open-source, plugin-based platform for multichannel electrophysiology. J. Neural Eng. 14, 045003 (2017).

48. Viejo, G., et al. Pynapple, a toolbox for data analysis in neuroscience. Preprint at 10.7554/eLife.85786.2 (2023).

49. Dong, Z. et al. Minian, an open-source miniscope analysis pipeline. eLife 11, e70661 (2022).

50. Chang, H. et al. Sleep microstructure organizes memory replay. Nature 637, 1161–1169 (2025).

51. Mölter, J., Avitan, L. & Goodhill, G. J. Detecting neural assemblies in calcium imaging data. BMC Biol 16, 143 (2018).

52. Lopes-dos-Santos, V., Ribeiro, S. & Tort, A. B. L. Detecting cell assemblies in large neuronal populations. Journal of Neuroscience Methods 220, 149–166 (2013).

53. Bates, D., Mächler, M., Bolker, B. & Walker, S. Fitting Linear Mixed-Effects Models Using lme4. J. Stat. Soft. 67, (2015).

